# Genome-wide identification of Arabidopsis non-AUG-initiated upstream ORFs with evolutionarily conserved regulatory sequences that control protein expression levels

**DOI:** 10.1101/2021.03.25.436978

**Authors:** Yuta Hiragori, Hiro Takahashi, Noriya Hayashi, Shun Sasaki, Kodai Nakao, Taichiro Motomura, Yui Yamashita, Satoshi Naito, Hitoshi Onouchi

## Abstract

Upstream open reading frames (uORFs) are short ORFs found in the 5′-UTRs of many eukaryotic transcripts and can influence the translation of protein-coding main ORFs (mORFs). Recent genome-wide ribosome profiling studies have revealed that thousands of uORFs initiate translation at non-AUG start codons. However, the physiological significance of these non-AUG uORFs has so far been demonstrated for only a few of them. It is conceivable that physiologically important non-AUG uORFs are evolutionarily conserved across species. In this study, using a combination of bioinformatics and experimental approaches, we searched the Arabidopsis genome for non-AUG-initiated uORFs with conserved sequences that control the expression of the mORF-encoded proteins. As a result, we identified four novel regulatory non-AUG uORFs. Among these, two exerted repressive effects on mORF expression in an amino acid sequence-dependent manner. These two non-AUG uORFs are likely to encode regulatory peptides that cause ribosome stalling, thereby enhancing their repressive effects. In contrast, one of the identified regulatory non-AUG uORFs promoted mORF expression by alleviating the inhibitory effect of a downstream AUG-initiated uORF. These findings provide insights into the mechanisms that enable non-AUG uORFs to play regulatory roles despite their low translation initiation efficiencies.

## INTRODUCTION

The 5′-UTRs of many eukaryotic transcripts contain potentially translatable short open reading frames, referred to as upstream ORFs (uORFs). Since 43S pre-initiation complexes (PICs) scan for a start codon along an mRNA from the 5′ end in eukaryotes, PICs may recognize the start codon of a uORF and translate the uORF before reaching the downstream main ORF (mORF). uORF translation affects the expression of mORF-encoded proteins in various ways. Most commonly, if a uORF is translated by a ribosome, the ribosome dissociates from the mRNA after terminating translation at the uORF stop codon (1, 2). In this case, the PICs captured by the uORF do not translate the downstream mORF, and therefore, the magnitude of the inhibitory effect of the uORF on mORF translation correlates with the translation initiation efficiency of the uORF. When a uORF is short, the small ribosomal subunit may remain bound to the mRNA after terminating translation of the uORF, then resume scanning and reinitiate translation at a downstream start codon (3). If the reinitiation efficiency at the mORF start codon is high, the uORF has only a small impact on mORF translation. The reinitiation efficiency depends on the time required for uORF translation and the distance between the uORF stop codon and the downstream start codon (4–7). The intercistronic distance needed for efficient reinitiation depends on the cellular availability of the ternary complex that comprises eukaryotic initiation factor 2 (eIF2), GTP, and Met-tRNA_i_^Met^, and the level of the available ternary complex decreases under various stress conditions (8). The stress-responsive translational regulation of yeast *GCN4* and mammalian *ATF4* and *ATF5* mRNAs relies on these properties of reinitiation (8–13). These mRNAs contain an inhibitory uORF between the 5′-most uORF and the mORF. Under non-stress conditions, after translating the 5′-most uORF, ribosomes preferentially reinitiate translation at the start codon of the inhibitory uORF, resulting in repression of mORF translation. In contrast, under stress conditions, reinitiation is delayed due to the reduced availability of the ternary complex. Therefore, ribosomes more frequently bypass the inhibitory uORF and reinitiate translation at the start codon of the mORF, resulting in enhanced mORF translation. Another major mechanism by which uORFs control mORF translation involves ribosome stalling caused by uORF-encoded peptides. In this mechanism, the nascent peptide encoded by a uORF acts inside the ribosomal exit tunnel to cause the ribosome to stall during translational elongation or termination (14). Ribosome stalling on the uORF results in translational repression of the downstream mORF because the stalled ribosome blocks the access of other scanning PICs to the mORF start codon (15). In some genes, uORF peptide-mediated ribosome stalling is involved in translational regulation in response to metabolites or environmental stresses (14, 16, 17). In these regulatory mechanisms, the uORF translation initiation efficiency or the ribosome stalling efficiency is regulated in a condition-dependent manner. In addition to the effects of uORFs on mORF translation, uORFs can affect mRNA stability through the nonsense-mediated mRNA decay (NMD) pathway. Many uORF-containing transcripts have been shown to be targeted by NMD (18). Additionally, it has been reported that, in some genes, ribosome stalling at the uORF stop codon promotes NMD (19, 20).

Although most uORFs that have been shown to play regulatory roles begin with a canonical AUG codon, several uORFs with non-AUG start codons have also been reported to regulate mORF translation (21–25). For example, the polyamine-responsive translational regulation of the mouse antizyme inhibitor (*AZIN1*) mRNA involves an AUU-initiated uORF (21). In this regulation, polyamine induces ribosome stalling at the Pro-Pro-Trp motif of the uORF by inhibiting the translation elongation factor eIF5A (26). The stalled ribosome results in ribosome queuing by blocking scanning PICs and other translating ribosomes. This ribosome queuing promotes translation initiation at the AUU start codon of the uORF by positioning a PIC near the uORF start codon, thereby repressing mORF translation. Since AZIN1 is a positive regulator of polyamine synthesis, this translational regulation functions as a part of a negative feedback loop of polyamine biosynthesis. In plants, feedback regulation of ascorbate biosynthesis involves translational regulation mediated by a non-AUG uORF. The *GGP* gene in kiwifruit codes for a GDP-L-galactose phosphorylase, a major control enzyme in the ascorbate biosynthesis pathway, and has an evolutionarily conserved ACG-initiated uORF in its 5′-UTR. This non-AUG uORF mediates ascorbate-responsive translational repression of the downstream mORF in an amino acid sequence-dependent manner (22).

As exemplified by the non-AUG uORFs in *AZIN1* and *GPP*, translation can be initiated at near-cognate codons (NCCs) that differ from the canonical AUG codon by a single nucleotide. Among NCCs, translation initiation at CUG codons is generally most efficient, although the reported initiation efficiencies at NCCs differ among studies depending on experimental systems (27, 28). In mammalian cells, the translation initiation efficiency at CUG is 20-40% of that at AUG when in an optimal sequence context (29, 30). The second most efficient NCC is GUG or ACG, with 7-20% efficiencies compared with AUG (29, 30). The other NCCs are less efficient. In particular, the initiation efficiencies of AAG and AGG are very low (27, 28). Similarly, an early study reported that, in plant protoplasts, the initiation efficiencies of CUG, GUG, and ACG were 30%, 15%, and 15%, respectively, whereas the other NCCs were less efficient (31). The translation initiation efficiencies at NCCs are more dependent on the sequence context than that of AUG (30). Therefore, the difference in the translation initiation efficiencies between AUG and NCCs are generally larger in non-optimal contexts.

The genome-wide identification of translated uORFs by recent ribosome profiling studies revealed that thousands of uORFs are translated from NCCs (32–34). However, to date, the physiological significance of these NCC-initiated uORFs has been demonstrated for only a few of them. Given that the efficiencies of translation initiation at NCCs are generally low, most of the individual NCC-initiated uORFs should have only small impacts on the expression of mORF-encoded proteins. One possible mechanism by which an NCC-initiated uORF may exert a substantial impact on mORF expression is that the nascent peptide encoded by the NCC-initiated uORF causes ribosome stalling. If an NCC-initiated uORF causes ribosome stalling when translated, the stalled ribosome blocks other scanning PICs, and therefore, the NCC-initiated uORF may strongly repress mORF translation despite its low translation initiation efficiency. Additionally, ribosome stalling on a uORF may induce mRNA degradation via NMD if ribosomes stall at the stop codon of the uORF (19, 20). In fact, the regulatory uORF of the *Neurospora crassa arg2* gene, one of the most well-characterized AUG-initiated regulatory uORFs, has a low translation initiation efficiency (15). This uORF causes ribosome stalling in response to arginine (Arg), thereby regulating translation of the mORF, which codes for a subunit of the key enzyme of Arg biosynthesis. The translation initiation efficiency of the *arg-2* uORF is thought to need to be low to allow efficient mORF translation under low-Arg conditions and to repress mORF translation only when ribosome stalling occurs (15).

Since most known regulatory uORFs that cause ribosome stalling encode evolutionarily conserved peptide sequences, NCC-initiated uORFs encoding conserved peptide sequences are likely to have physiologically important regulatory roles. Recently, van der Horst *et al*. searched the Arabidopsis genome for conserved peptide-encoding uORFs (CPuORFs), including those lacking AUG start codons, by comparing the 5′-UTR sequences between Arabidopsis and 31 other plant species, and 14 non-AUG CPuORFs were identified (35). In yeasts, 788 conserved non-AUG uORFs, including those whose sequences or positions are conserved among *Saccharomyces* species, have recently been identified using machine-learning to analyze ribosome profiling data and a comparative genomic approach (36). Examination of the effects of five of these conserved non-AUG uORFs using reporter assays revealed that the presence of four of these uORFs affects mORF expression (36). However, the effects of the conserved sequences of these non-AUG uORFs identified by these studies on mORF expression remain to be experimentally validated.

Here, we report the genome-wide identification of Arabidopsis regulatory non-AUG uORFs with evolutionarily conserved sequences by a combination of bioinformatics and experimental approaches. We searched for Arabidopsis NCC-initiated uORFs whose NCCs and encoded sequences were conserved across more than four orders and tested their effects on mORF expression using transient expression assays. As a result, three inhibitory non-AUG uORFs were identified. Among these, two repress mORF expression in a sequence-dependent manner. In addition, we identified one non-AUG uORF that promoted mORF expression by alleviating the inhibitory effect of a downstream AUG-initiated uORF. We discuss the potential regulatory roles of these evolutionarily conserved non-AUG uORFs and the underlying mechanisms that enable non-AUG uORFs to play regulatory roles despite their low translation initiation efficiencies.

## MATERIALS AND METHODS

### Extraction of NCC-initiated uORF sequences

To extract NCC-initiated uORF sequences from the Arabidopsis (*Arabidopsis thaliana*) genome, we used genome sequence files in FASTA format and genomic coordinate files in GFF3 format, obtained from Ensembl Plants Release 33 (https://plants.ensembl.org/index.html) (37). Based on genomic coordinate information, we extracted exon sequences from the Arabidopsis genome sequence and constructed a transcript sequence dataset by combining exon sequences. We extracted 5′-UTR sequences from the transcript sequence dataset on the basis of the transcription start site and the translation start codon of each transcript in the genomic coordinate files,. Then, we searched the 5′-UTR sequences for NCCs other than AAG and AGG. When there were multiple NCCs (other than AAG and AGG) in the same reading frame, the 5′-most NCC was selected. If there was an in-frame AUG codon upstream of the 5′-most NCC, none of the NCCs in the reading frame was selected. The sequence starting with the selected NCC and ending with the nearest downstream in-frame stop codon were extracted as an NCC-initiated uORF sequence.

### Calculation of uORF-mORF fusion ratio

The uORF-mORF fusion ratio for each of the extracted NCC-initiated uORFs was calculated as previously described (38).

### BLAST-based search for uORFs conserved between homologous genes

uORF-tBLASTn and mORF-tBLASTn analyses were performed as previously described (38), with some modifications. The plant (Viridiplantae) transcript sequence database previously described (38) was used for the uORF-tBLASTn and mORF-tBLASTn analyses. This transcript sequence database comprised the sequences of the National Center for Biotechnology Information (NCBI) Reference Sequence (RefSeq) transcripts, contigs of assembled expressed sequence tags (ESTs) and transcriptome shotgun assemblies (TSAs), and unclustered singleton ESTs and TSAs. In the uORF-tBLASTn analysis, the amino acid sequence of each of the extracted Arabidopsis NCC-initiated uORFs was queried against the plant transcript sequence database, using NCBI tBLASTn with an *E*-value threshold of 2000. For each of the uORF-tBLASTn hits, the downstream in-frame stop codon closest to the 5′-end of the matching region was selected (Supplementary Figure S1). Then, an in-frame ATG codon or NCC other than AAG and AGG was searched upstream of the selected stop codon. uORF-tBLASTn hits containing neither ATG codon nor NCC upstream of the selected stop codon were discarded. If one or more in-frame ATG codons or NCCs were identified, the 5′-most ATG codon or NCC was selected (Supplementary Figure S1). The ORF beginning with the selected ATG codon or NCC and ending with the selected stop codon was extracted as a putative uORF. In the mORF-tBLASTn analysis, the amino acid sequence of the mORF in the original Arabidopsis NCC uORF-containing transcript was queried against the downstream sequences of the putative uORFs, using tBLASTn with an *E*-value cutoff of 10^−1^. For each of the mORF-tBLASTn hits, the upstream in-frame stop codon closest to the 5′-end of the region matching the original mORF was selected, and the 5′-most in-frame ATG codon located downstream of the selected stop codon was identified as the putative mORF start codon (Supplementary Figure S1). If the putative mORF start codon was found downstream of the stop codon of the putative uORF, the mORF-tBLASTn hit sequence was extracted as a uORF-mORF homolog. Potential contaminant sequences derived from contaminating organisms, such as parasites, plant-feeding insects and infectious microorganisms, were excluded from the uORF-mORF homologs, as previously described (38). Based on taxonomic lineage information of EST, TSA, and RefSeq transcript sequences, provided by NCBI Taxonomy (https://www.ncbi.nlm.nih.gov/taxonomy), the uORF-mORF homologs were classified into orders and the 13 taxonomic categories previously defined (i.e. lamiids, asterids other than lamiids, mavids, fabids, eudicots other than rosids and asterids, commelinids, monocots other than commelinids, Angiospermae other than eudicots and monocots, Gymnospermae, Polypodiopsida, Embryophyta other than Euphyllophyta, Streptophyta other than Embryophyta, and Viridiplantae other than Streptophyta) (38). Arabidopsis NCC-initiated uORFs were selected as candidate non-AUG CPuORFs if their uORF-mORF homologs were found in fabids and at least four orders other than Brassicales.

### *K*_a_/*K*_s_ analysis

To generate the multiple uORF sequence alignment and to calculate the *K*_a_/*K*_s_ ratio for each candidate non-AUG CPuORF, one representative uORF-mORF homolog was selected from each order in which uORF-mORF homologs were identified, using the criteria described in our previous report (38). Then, the uORF sequence beginning with the conserved NCC or a different codon at the corresponding position was extracted from each representative uORF-mORF homolog. Only uORF sequences in representative uORF-mORF homologs selected from orders belonging to Angiospermae were used for *K*_a_/*K*_s_ analysis. Multiple alignments of the uORF amino acid sequences were generated by using standalone Clustal Omega version 1.2.2 (39) with the default parameters. For each candidate CPuORF, the median *K*_a_/*K*_s_ ratio for all pairwise combinations of the original Arabidopsis uORF and its homologous uORFs was calculated as previously described (38), using the kaks function in the seqinR package version 3.4.5 (40).

### Assessment of conservation of NCCs and their initiation context types

For each of the candidate non-AUG CPuORFs, NCCs and their initiation context types were manually compared among the original Arabidopsis uORF and its homologous uORFs in the representative uORF-mORF homologs, using the original uORF-containing transcript sequence and the transcript sequences of the uORF-mORF homologs.

### Assessment of overlap between candidate non-AUG CPuORFs and unknown conserved AUG uORFs

In the seventh step of our search for conserved NCC-initiated uORFs, we manually checked whether each candidate non-AUG CPuORF overlaps with an unknown conserved AUG uORF. A candidate was discarded if most of its representative uORF-mORF homologs contained a conserved ATG codon in any reading frame upstream of the conserved NCCs, without any stop codon between the conserved ATG codon and the conserved NCC in the same reading frame as the conserved ATG codon. Additionally, a candidate was discarded if the NCC-initiated uORFs in most of its representative uORF-mORF homologs contained a conserved ATG codon in any reading frame and the distance between the conserved NCC and the conserved ATG codon was less than 21 nt.

### Plasmid construction

To clone the 5′-UTRs of the Arabidopsis genes used for the transient expression assays, total RNA was extracted from one-week-old *A. thaliana* (Col-0 ecotype) seedlings using the RNeasy Mini kit (Qiagen), and cDNA was prepared from the total RNA using SuperScript III Reverse Transcriptase (Thermo Fisher Scientific) and an oligo(dT) primer (Life Technologies). The 5′-UTRs of each of the Arabidopsis genes were amplified by PCR from the cDNA using primers listed in Supplementary Tables S1 and S2. To construct reporter plasmids, the amplified 5′-UTR fragments were inserted between the *Xba*I and *Sal*I sites of plasmid pKM56 (41), which harbored the cauliflower mosaic virus 35S RNA (35S) promoter, the coding sequence of a Brazilian click beetle luciferase (Emerald luciferase: Eluc) with a PEST degradation sequence (Eluc-PEST), and the polyadenylation signal of the *A. thaliana HSP18.2* gene in pUC19, using the SLiCE method (42). The 5′-UTR DNA fragments of the *Theobroma cacao APD6* and *MYB7* orthologs and the *Arachis ipaensis FH11* ortholog were synthesized by Eurofins Genomics KK, based on NCBI RefSeq transcripts (GenBank accession numbers: XM_007041714.2, XM_007041801.2, and XM_016321042.2). The synthesized 5′-UTR fragments were individually ligated between the *Xba*I and *Sal*I sites of pKM56. Mutations were introduced into each of the 5′-UTRs using overlap extension PCR (43), with primers listed in Supplementary Tables S1 and S2. Sequence analysis confirmed the integrity of the PCR-amplified regions of all constructs.

### Transient expression assay

To prepare protoplasts, MM2d suspension cells (44) were collected by centrifugation on the third day after transfer to fresh media and were suspended in modified Linsmaier and Skoog (LS) medium (45) containing 1% (w/v) cellulase Onozuka RS (Yakult Pharmaceutical Industry), 0.5% (w/v) pectolyase Y23 (Seishin Pharmaceutical), and 0.4 M mannitol. Then, the cells were incubated at 26°C with gentle shaking until the suspension became turbid with protoplasts. The protoplasts were washed five times with wash buffer (0.4 M mannitol, 5 mM CaCl_2_, and 0.5 M 2-(*N*-morpholino)ethanesulfonic acid, pH 5.8) and were suspended in MaMg solution (5 mM morpholinoethanesulfonic acid, 15 mM MgCl_2_, and 0.4 M mannitol, pH 5.8). Then, 6 μl of DNA solution containing 3 μg each of the reporter plasmid and the *35S::Rluc* internal control plasmid (pKM5) (41), which contained the *Renilla* luciferase (Rluc) coding sequence under the control of the 35S promoter, were mixed with 100 μl of MaMg solution containing 1.5 × 10^5^ protoplasts and 106 μl of polyethylene glycol (PEG) solution (40% PEG4000, 0.1 mM CaCl_2_, and 0.2 M mannitol). This mixture was incubated for 15 min at room temperature and was diluted by adding 800 μl of wash buffer. The protoplasts were centrifuged and resuspended in 1 ml of the modified LS medium containing 0.4 M mannitol. After 24 h of incubation at 22°C in the dark, cells were harvested and disrupted in 100 or 200 μl of extraction buffer [100 mM(NaH_2_/Na_2_H)PO_4_ and 5 mM DTT, pH 7] by voltexing for 2 min. A Dual-Luciferase Reporter Assay kit (Promega) was used to measure the Rluc and Eluc activities.

## RESULTS

### Identification of Arabidopsis NCC-initiated uORFs with conserved sequences

Recently, we developed the ESUCA pipeline to efficiently perform genome-wide searches for AUG-initiated CPuORFs (38). ESUCA compares uORF amino acid sequences between certain species and any other species with available transcript databases. Therefore, ESUCA can be used to comprehensively identify CPuORFs conserved across various taxonomic ranges. In the present study, we modified ESUCA to search for NCC-initiated CPuORFs in the Arabidopsis genome. We first extracted ORFs beginning with an NCC other than AAG and AGG (i.e. CTG, GTG, TTG, ACG, ATA, ATT, or ATC) from the 5′-UTRs of Arabidopsis transcript sequences (Figure 1, step 1). Since many previous studies have shown that the efficiency of translation initiation at AGG and AAG are close to the background level (27, 28, 30), we did not extract ORFs starting with AAG or AGG. When there were multiple NCCs in the same reading frame, only the ORF beginning with the 5′-most NCC was extracted. However, we did not extract the ORF if an in-frame AUG codon precedes the 5′-most NCC. We also did not extract ORFs starting with an NCC in the 5′-UTR of a gene and ending with a stop codon in the mORF. Although an ORF that starts with an start codon in the 5′-UTR and overlaps with the downstream mORF is also usually considered a uORF, we focused on NCC-initiated uORFs that have both start and stop codons within the 5′-UTR because uORF sequences overlapping with the mORFs may be conserved due to functional constraints of the mORF-encoded proteins. We also removed NCC-initiated uORFs overlapping the AUG-initiated CPuORFs previously identified by ESUCA (38) to exclude those whose sequences were conserved due to functional constraints of the overlapping AUG-initiated CPuORFs (Figure 1, step 2).

**Figure 1.**
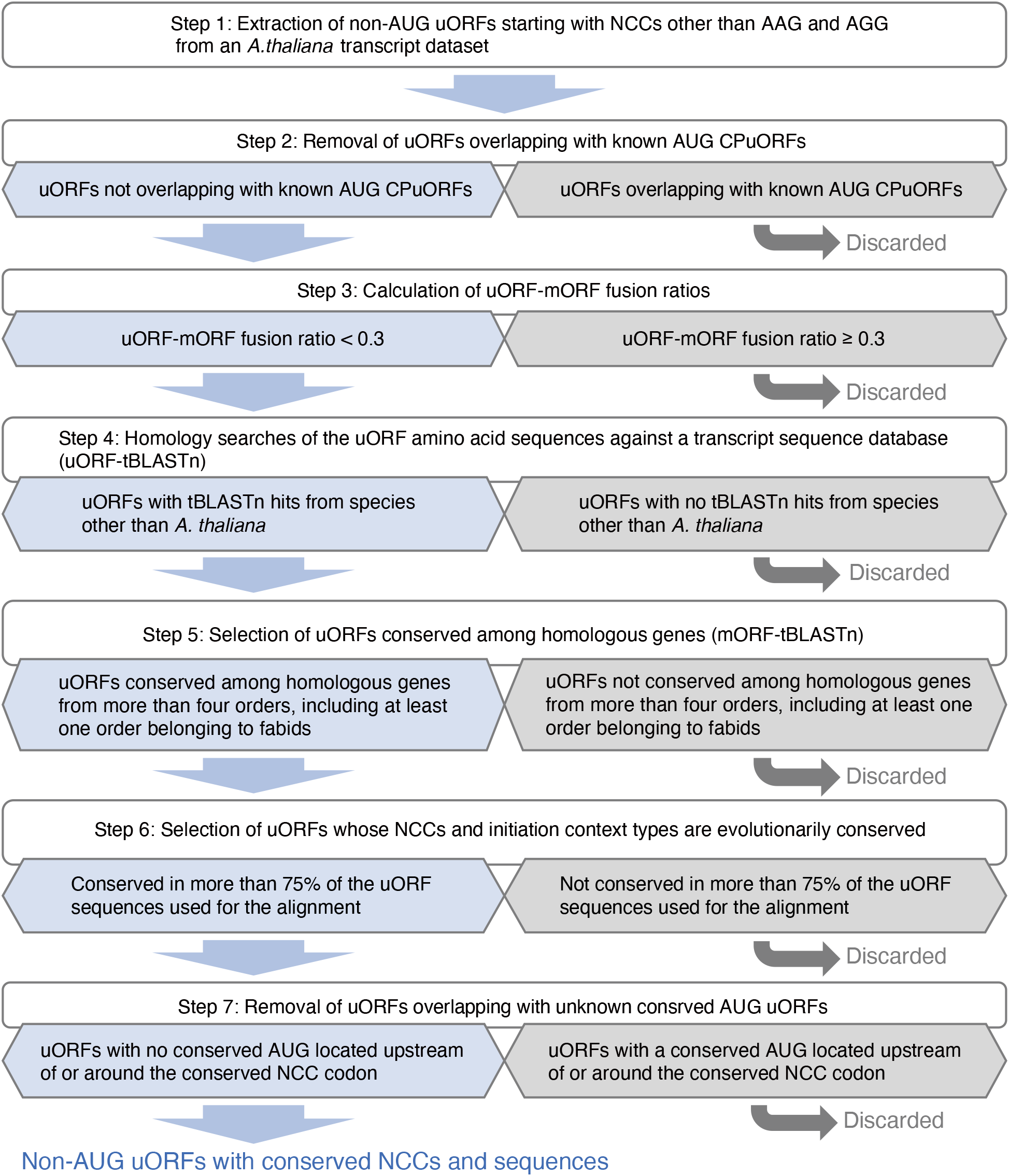
Outline of the search for Arabidopsis non-AUG uORFs with conserved NCCs and sequences.

We extracted NCC-initiated uORFs from all the splice variants of the Arabidopsis genes in the Ensembl Plants transcript database (http://plants.ensembl.org/index.html) (37). In our previous genome-wide searches for AUG-initiated CPuORFs, there were cases where a uORF found in the 5′-UTR of a splice variant was fused to the mORF in another splice variant from the same gene (38, 46). In some of these cases, the uORF sequence was found to be conserved because it encoded a part of the mORF-encoded protein sequence (38, 46). To distinguish between such ‘spurious’ CPuORFs and ‘true’ CPuORFs, which are evolutionarily conserved because of functional constraints of their encoded small peptides, we recently developed an algorithm to assess whether transcripts bearing a uORF-mORF fusion are minor or major forms among orthologous transcripts (38). For each uORF, this algorithm calculates the ratio of the NCBI RefSeq transcripts with a uORF-mORF fusion to all RefSeq transcripts with both sequences similar to the uORF and its downstream mORF. This ratio was termed the uORF-mORF fusion ratio (38). We calculated the uORF-mORF fusion ratios of the NCC-initiated uORFs extracted in step 2 and discarded those with uORF-mORF fusion ratios equal to or greater than 0.3 (Figure 1, step 3). We also removed uORFs whose numbers of RefSeq transcripts containing both sequences similar to the uORF and its downstream mORF were less than 10. This was done because the appropriate evaluation of uORF-mORF fusion ratios was difficult with a few related RefSeq transcripts, and such uORF sequences are unlikely to be evolutionarily conserved.

Next, we performed homology searches of the uORF amino acid sequences, using algorithms in the ESUCA pipeline with some modifications. In this search, each of the NCC-initiated uORF amino acid sequences was queried against a plant transcript sequence database that contained the sequences of RefSeq transcripts, assembled EST/TSA contigs, and unclustered singleton ESTs and TSAs, using tBLASTn with an *E*-value cutoff of 2000 (uORF-tBLASTn analysis) (Figure 1, step 4). For each uORF-tBLASTn hit, the downstream in-frame stop codon closest to the 5′-end of the matching region was selected. Then, we looked for an in-frame ATG codon or NCC other than AAG and AGG (i.e. ATG, CTG, GTG, TTG, ACG, ATA, ATT, or ATC) upstream of the selected stop codon and the 5′-most ATG codon or NCC was selected (Supplementary Figure S1). The ORF beginning with the selected ATG codon or NCC and ending with the selected stop codon was extracted as a putative uORF. The downstream sequences of the putative uORFs were subjected to another tBLASTn search to determine whether the uORF-tBLASTn hits were derived from homologs of the original Arabidopsis uORF-containing transcript (Supplementary Figure S1). In this analysis, the amino acid sequence of the mORF in each original uORF-containing transcript was used as a query, and the uORF-tBLASTn hits matching the mORF with an *E*-value less than 10^-1^ were extracted (mORF-tBLASTn analysis) (Figure 1, step 5). If a uORF-tBLASTn hit contained a partial or intact ORF sequence similar to the original mORF sequence downstream of the putative uORF, it was selected as a uORF-mORF homolog of the original Arabidopsis uORF-containing transcript.

Our previous study on the relationship between the taxonomic range of CPuORF sequence conservation and their sequence-dependent effects on mORF translation showed that AUG-initiated CPuORFs conserved only among rosids can have sequence-dependent regulatory effects (38). Rosids consists of malvids, to which Arabidopsis belongs, and fabids. Therefore, to identify non-AUG uORFs conserved in a wide range of rosids, we selected Arabodppsis NCC-initiated uORFs for which uORF-mORF homologs were found in fabids. Among these Arabidopsis NCC-initiated uORFs, we further selected those for which uORF-mORF homologs were found in at least four orders other than Brassicales, to which Arabidopsis belongs, to compare the start codons and their sequence contexts among representative uORFs from more than four orders at a later step. According to these criteria, 108 Arabodppsis NCC-initiated uORFs were selected as candidate non-AUG CPuORFs.

For each candidate non-AUG CPuORF, one representative uORF-mORF homolog was selected from each order in which uORF-mORF homologs were identified, and the putative uORF sequences in the selected uORF-mORF homologs and the original Arabidopsis NCC-initiated uORF sequence were used to generate multiple amino acid and nucleotide sequence alignments. If the amino acid sequence of a uORF is evolutionarily conserved because of functional constraints of the uORF-encoded peptide, the amino acid sequence in the functionally important region of the peptide is expected to be conserved among the uORF and its orthologous uORFs. Therefore, we manually checked whether the amino acid sequences in the same region were conserved among the uORF sequences in the alignment of each candidate non-AUG CPuORF. Then, we removed sequences that did not share the consensus amino acid sequence in the conserved region. When this change resulted in the number of orders with the uORF-mORF homologs becoming less than four, the candidate non-AUG CPuORFs were discarded. In addition, when multiple candidate non-AUG CPuORFs derived from splice variants of the same gene partially or completely shared amino acid sequences, the one with the longest conserved region was manually selected based on the uORF amino acid sequence alignments. When multiple candidate non-AUG CPuORFs overlapped with each other in different frames, the one conserved in the greatest number of orders was retained and the others were discarded.

As mentioned in the Introduction, translation initiation efficiencies differ among different NCCs (27, 28). Additionally, sequence contexts surrounding NCCs affect the efficiencies of translation initiation at the NCCs. In particular, nucleotides at the ‒3 and +4 positions, where the first nucleotide of an NCC is +1, are critical for the translation initiation efficiencies at NCCs. Although a purine (A or G) at −3 and a G at +4 are favorable for efficient translation initiation of both AUG scodons and NCCs, translation initiation at NCCs is more dependent on this sequence context than that at AUG codons, and the effects of G at +4 are different among NCCs (30). Therefore, if a non-AUG uORF is conserved because of its regulatory role, it is likely that the non-AUG start codon and its sequence context are evolutionarily conserved. We classified the sequence contexts of NCCs in the remaining candidate non-AUG CPuORFs into four types: (1) optimal context; (2) ‒3 sub-optimal context; (3) +4 sub-optimal context; (4) non-optimal context. The optimal context contains both a purine at ‒3 and a G at +4. The ‒3 sub-optimal context contains a purine at ‒3, but does not have a G at +4. The +4 sub-optimal context contains a G at +4, but does not have a purine at ‒3. The non-optimal context contains neither a purine at −3 nor a G at +4. The candidate non-AUG CPuORFs identified by our search included the known regulatory non-AUG CPuORFs of the Arabidopsis *VTC2* and *VTC5* genes, which are orthologs of the kiwifruit *GGP* gene (22, 47). In 90% and 86% of the *VTC2* and *VTC5* orthologous transcript sequences used for the alignments, respectively, the ACG start codons of the uORFs were conserved and their initiation contexts were optimal. Therefore, we selected candidate non-AUG CPuORFs for which more than 75% of sequences used for the uORF sequence alignment have conserved NCCs with the same initiation context type upstream of the sixth codon of the conserved region (Figure 1, step 6). Using these criteria, 19 candidate non-AUG CPuORFs were selected. For seven of the 19 candidates, conserved AUG codons were found upstream of or close to the conserved NCCs. Therefore, the sequences of these seven candidates may be conserved due to functional constraints of the overlapping uORFs starting with the conserved AUG codons rather than functional constraints of the candidate non-AUG CPuORFs. For this reason, we discarded these seven candidates. If the conserved AUG codon of an overlapping AUG uORF was more than 20 nt downstream of the conserved NCCs of the candidate CPuORFs, we did not discard the candidates (Figure 1, step 7). As a result, 12 non-AUG uORFs with conserved NCCs and sequences were identified (Table 1, Figure 2, Supplementary Figure S2, Supplementary Table S3).

**Table 1.** Novel and known conserved Arabidopsis non-AUG uORFs identified in this study.

| Gene<br>(AGI code) | Gene<br>symbol | Conserved NCC | Initiation<br>context type | *K<sub>a</sub>/K<sub>s</sub>* analysis of uORF sequences |  | Novel or<br>known |
| --- | --- | --- | --- | --- | --- | --- |
|  |  |  |  | Median pairwise<br>*K<sub>a</sub>/K<sub>s</sub>* ratio | *U*-test<br>*p value* |  |
| AT1G32240 | *KAN2* | GUG | Non-optimal | 0.63 | 2.9E-20 | Novel |
| AT1G53300 | *TTL1* | AUU | Non-optimal | 0.59 | 4.7 E-04 | Novel |
| AT1G58290 | *HEMA1* | GUG | Optimal | 0.47 | 6.4E-05 | Novel |
| AT2G16720 | *MYB7* | CUG (GUG in Arabidopsis) | -3 sub-optimal | 0.44 | 3.8E-02 | Novel |
| AT2G20080 | *SPEAR1* | AUC | -3 sub-optimal | 0.30 | 8.0E-02 | Novel |
| AT3G05470 | *FH11* | CUG (UUG in Arabidopsis) | Non-optimal | 0.44 | 2.1E-06 | Novel |
| AT3G08850 | *RAPTOR1B* | CUG | Optimal | 0.14 | 5.7E-89 | Known |
| AT4G03260 | *MASP1* | CUG | Optimal | 0.31 | 1.4E-259 | Known |
| AT4G19110 | *MPK22* | CUG | Optimal | 0.91 | 7.0E-01 | Novel |
| AT4G26850 | *VTC2* | ACG | Optimal | 0.22 | 0.0E+00 | Known |
| AT4G34710 | *SPE2* | CUG | Non-optimal | 0.91 | 4.6E-01 | Novel |
| AT4G38520 | *APD6* | UUG (GUG in Arabidopsis) | -3 sub-optimal | 0.75 | 3.5E-01 | Novel |
| AT5G35750 | *AHK2* | CUG | -3 sub-optimal | 0.60 | 6.3 E-02 | Novel |
| AT5G55120 | *VTC5* | ACG | Optimal | 0.22 | 0.0E+00 | Known |

**Figure 2.**
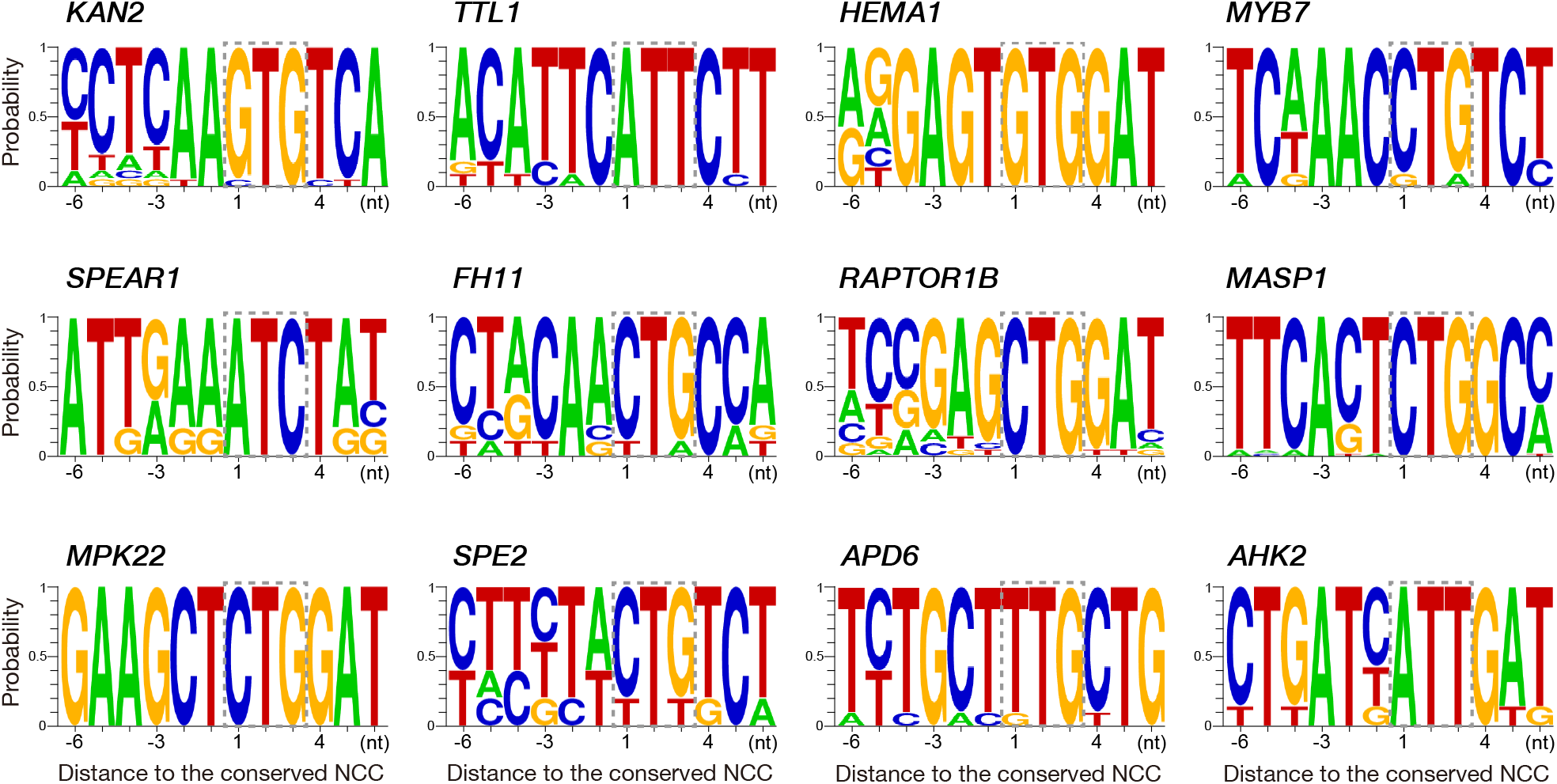
Sequence context of the conserved NCC codons of the non-AUG uORFs identified in this study. The conserved NCCs are surrounded by dotted lines. Logos were generated using WebLogo 2.8.2 (https://weblogo.berkeley.edu/logo.cgi). For each logo, the sequences surrounding the conserved NCCs of the original Arabidopsis uORF-containing transcript and its representative uORF-mORF homologs were used.

The 12 conserved non-AUG uORFs were subjected to calculations of *K*_a_/*K*_s_ ratios to assess whether the sequences of these non-AUG CPuORFs were conserved at the nucleotide or amino acid level. A *K*_a_/*K*_s_ ratio close to 1 indicates neutral evolution, whereas a *K*_a_/*K*_s_ ratio close to 0 suggests that purifying selection acted on the amino acid sequences (48). The *K*_a_/*K*_s_ ratios of five of the conserved non-AUG uORFs were less than 0.5 and significantly different from those of the negative controls, with *p* values less than 0.05 (Table 1), suggesting that the sequences of at least these five non-AUG uORFs were conserved at the amino acid level. Of these, the non-AUG uORFs of the *MASP1* and *RAPTOR1B* genes have recently been reported as non-AUG CPuORFs (35). Therefore, our search identified three novel non-AUG CPuORFs, namely those of the *HEMA1*, *MYB7*, and *FH11* genes.

### Effects of the conserved non-AUG uORFs on mORF expression

Next, we examined the effects of the conserved non-AUG uORFs on mORF expression using transient expression assays. Although the primary aim of this study was to identify regulatory non-AUG CPuORFs, we tested the effects of the 12 conserved non-AUG uORFs, irrespective of their *K*_a_/*K*_s_ ratios. This is because non-AUG uORFs with conserved start codons and conserved nucleotide sequences may have regulatory roles even if their sequences are not conserved at the amino acid level. In addition, investigating the effects of these non-AUG uORFs may allow us to gain insights into the differences in the regulatory effects and roles of amino acid sequence-dependent and -independent regulatory non-AUG uORFs.

The 5′-UTR sequences of the 12 genes bearing the conserved non-AUG uORFs were fused to the *Eluc-PEST* coding sequence and were placed under the control of the 35S promoter to generate the wild-type (WT) reporter constructs (Figures 3 and 4A, Supplementary Figure S3). To determine the effects of the conserved non-AUG uORFs on expression from the mORF, the conserved NCC in each reporter construct was mutated. In these NCC-mutant constructs, the second and/or third letter(s) of each conserved NCC was altered so that at least two nucleotides differed from ATG. The NCCs of the conserved uORFs of the Arabidopsis *MYB7*, *FH11*, and *APD6* genes are different from those of their orthologous uORFs (Table 1, Figure 2). For example, the conserved NCCs of the uORFs of *APD6* orthologous genes from orders other than Brassicales are UUG, whereas the corresponding NCC of the Arabidopsis *APD6* uORF is GUG (Figure 2, Supplementary Figure S3K). Therefore, for each of these three uORFs (*MYB7*, *FH11*, and *APD6* uORFs), we also created a reporter construct with the 5′-UTR of one of its orthologous genes (i.e. *Theobroma cacao MYB7* ortholog, *Arachis ipaensis FH11* ortholog, and *T. cacao APD6* ortholog) (Supplementary Figure S3M, N and O).

**Figure 3.**
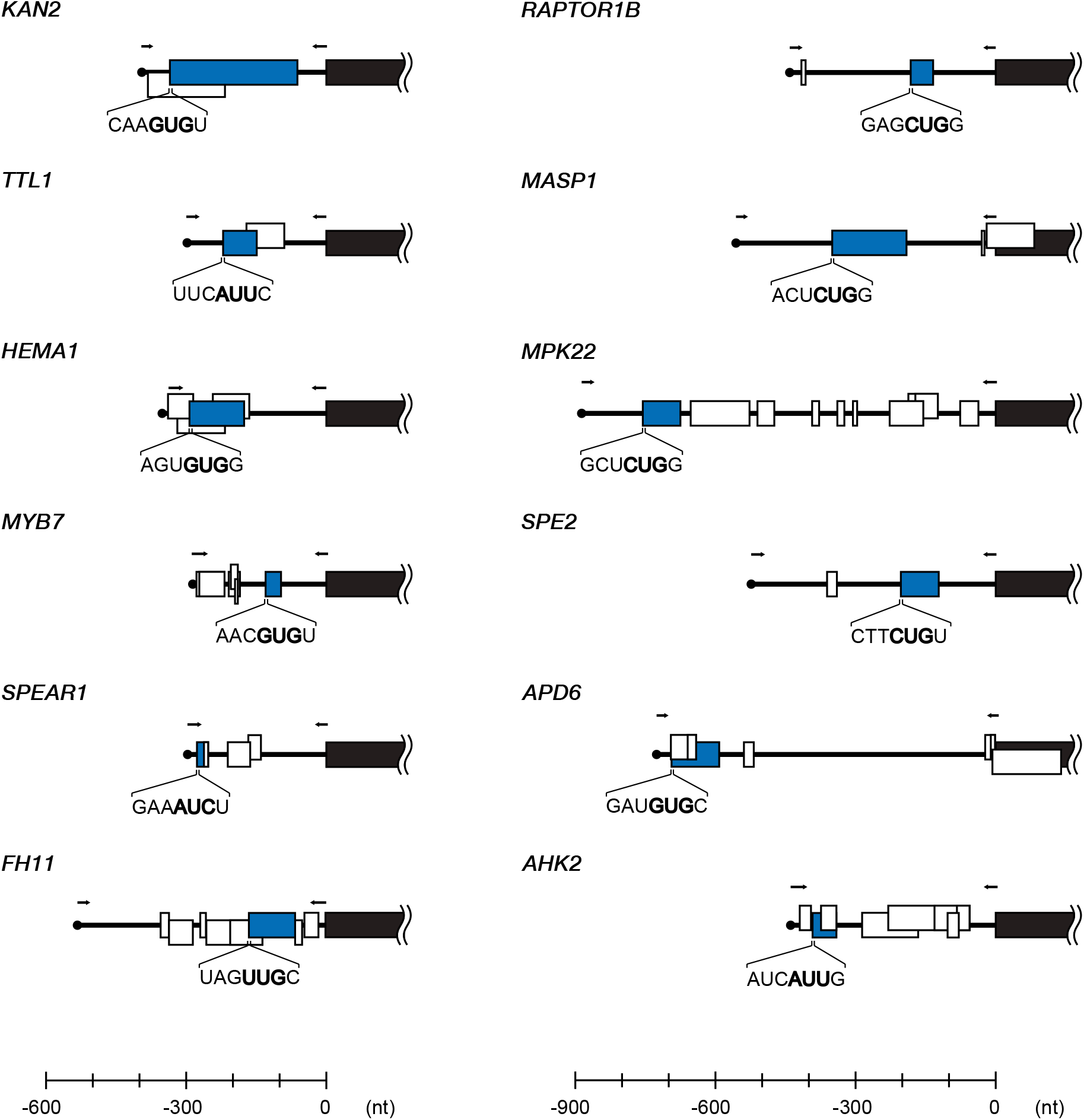
Schematic representation of the 5′-UTRs containing the conserved non-AUG uORFs identified in this study. Blue, white, and black boxes represent the conserved non-AUG uORFs, AUG-initiated uORFs, and the 5′-terminal parts of the mORFs, respectively. The translation initiation contexts of the NCCs of the conserved non-AUG uORFs are shown. The arrows indicate the position of the primers used to clone the 5′-UTRs for transient expression assay.

**Figure 4.**
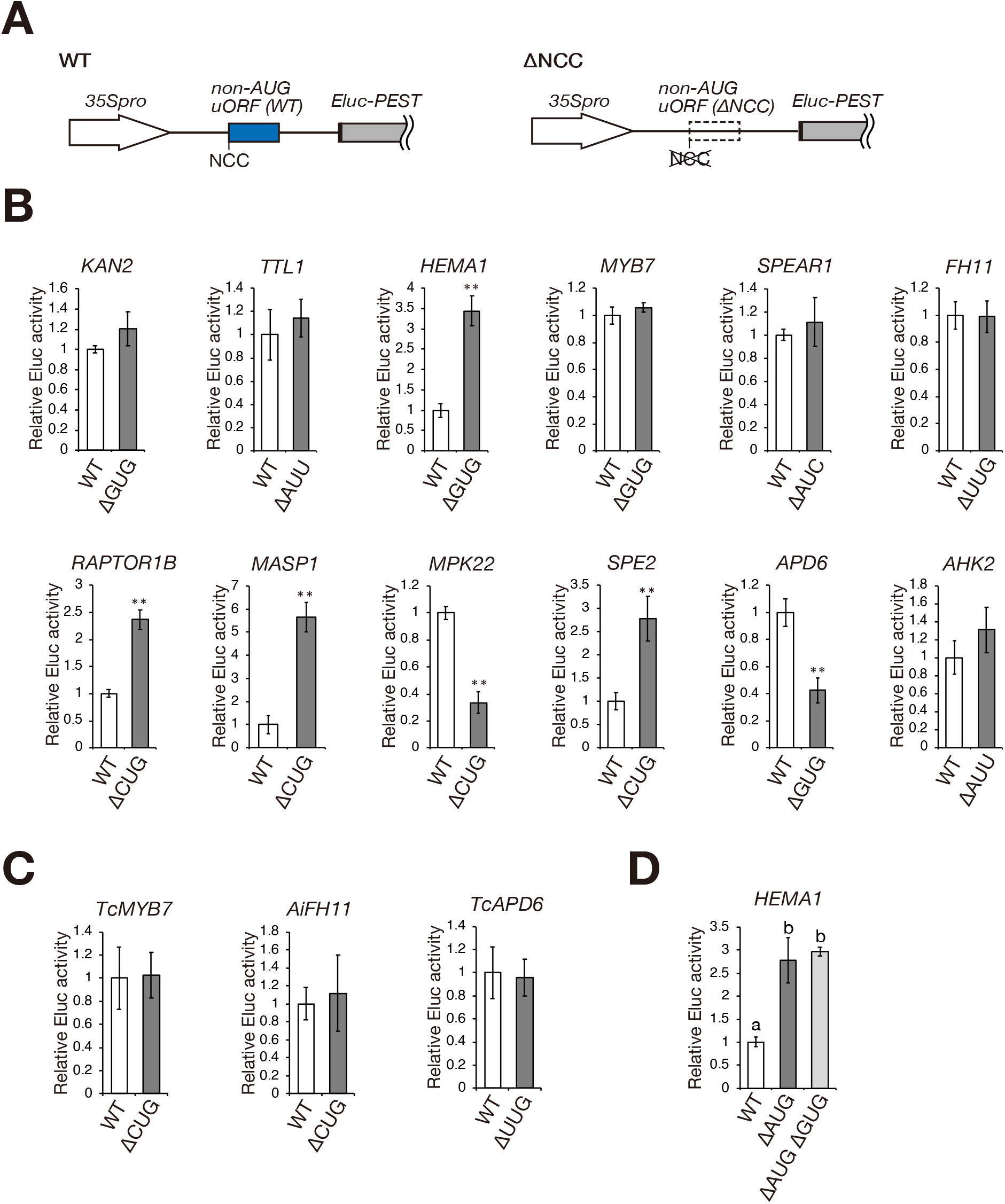
Effects of the presence of the conserved non-AUG uORFs on mORF expression. (A) Schematic representation of the wild-type (WT) and NCC mutant (ΔNCC) reporter constructs. The 5′-UTR containing each conserved non-AUG uORF was inserted between the 35S promoter (*35Spro*) and the *Eluc-PEST* coding sequence. The blue box in the WT reporter construct represents the conserved non-AUG uORF. The open box with a dotted line in the ΔNCC reporter construct depicts the NCC-mutant non-AUG uORF lacking the conserved NCC. The black boxes represent the first several nucleotides of the mORF. See Figure 3 and Supplementary Figure S3 for the exact position and length of each conserved non-AUG uORF and the 5′-UTR sequences used. (B, C) Transient expression assay. Each reporter plasmid containing the WT or mutant version of the 5′-UTR of the indicated gene was co-transfected into MM2d protoplasts with the *35S::Rluc* internal control plasmid Eluc activity was normalized to Rluc activity. Data represent the mean ± SD of the normalized Eluc activities of three biological replicates, relative to that of the corresponding WT reporter construct. Double asterisks indicate significant differences between the WT and mutant constructs at *P* < 0.01, respectively, as determined by Student’s *t*-test.

The reporter constructs were individually transfected into protoplasts from *Arabidopsis thaliana* MM2d suspension-cultured cells. After 24 h of incubation, the cells were harvested and disrupted to measure luciferase activity. Of the 15 conserved non-AUG uORFs tested, the NCC mutations of the *HEMA1*, *RAPTOR1B*, *MASP1*, and *SPE2* uORFs significantly increased reporter gene expression, whereas those of the *APD6* and *MPK22* uORFs significantly reduced reporter gene expression (Figure 4B and C). These results suggest that translation initiation occurs at these conserved NCCs and that the conserved non-AUG uORFs of *HEMA1*, *RAPTOR1B*, *MASP1*, and *SPE2* have repressive effects on mORF expression, whereas those of *APD6* and *MPK22* have positive effects. However, in the *APD6* uORF mutant, the third letter of the GUG codon was changed to C, and an AUG codon overlaps with the GUG codon two nucleotides upstream of the mutated site (Figure 3, Supplementary Figure S3K). Therefore, this mutation may affect the efficiency of translation initiation at the AUG codon, and the increased reporter gene expression may be due to the change in the translation initiation efficiency at the AUG codon rather than abolition of translation initiation at the GUG codon. In addition, mutating the conserved UUG codon of the non-AUG uORF of the *T. cacao APD6* ortholog did not significantly affect reporter gene expression (Figure 4C). These results suggest that the *APD6* non-AUG uORF does not have a regulatory effect on mORF expression. The *HEMA1* conserved non-AUG uORF also overlaps with AUG uORFs (Figure 3, Supplementary Figure S3C). Therefore, we tested the effect of the *HEMA1* non-AUG uORF on mORF expression in the absence of the upstream overlapping AUG uORFs. As shown in Figure 4D, mutating the conserved CUG codon of the *HEMA1* non-AUG uORF did not significantly affect reporter gene expression in the absence of the upstream overlapping AUG uORFs. This result suggests that the effect of the CTG-to-CTC mutation of the *HEMA1* non-AUG uORF was due to the change in the sequence of the overlapping AUG uORF rather than abolition of translation initiation at the conserved CUG codon, and that the *HEMA1* conserved non-AUG uORF does not have a regulatory effect on mORF expression. Therefore, our transient assay identified three conserved repressive non-AUG uORFs (the *RAPTOR1B*, *MASP1*, and *SPE2* uORFs) and one conserved promotive non-AUG uORF (the *MPK22* uORF).

### Identification of peptide sequence-dependent regulatory non-AUG uORFs

Several regulatory AUG uORFs with evolutionarily conserved amino acid sequences are known to encode peptides that repress mORF translation by causing ribosome stalling (15, 20, 41, 49–51). To determine whether the repressive effects of the *RAPTOR1B*, *MASP1*, and *SPE2* conserved non-AUG uORFs depend on their encoded peptide sequences, we altered the amino acid sequences of the WT and NCC mutant versions of these uORFs by introducing frameshift mutations. To change the amino acid sequences of the conserved regions of these uORFs, a +1 or −1 frameshift was introduced into the conserved region of each uORF, and another frameshift was introduced before the stop codon to shift the reading frame back to the original frame (Supplementary Figure S3G, H and J). In the frameshift mutant of the *SPE2* non-AUG uORF, an additional nucleotide substitution was introduced to remove an early in-frame stop codon in the shifted reading frame (Supplementary Figure S3J). For all the three non-AUG uORFs tested, the frameshift mutations introduced into the WT uORFs significantly elevated the expression of the downstream reporter gene (Figure 5B and C). For *RAPTOR1B* and *MASP1*, the frameshift mutations introduced into the NCC mutant uORFs caused no further significant increase in reporter gene expression (Figure 5B), indicating that the effects of the frameshift mutations depended on translation from the conserved CUG codons. These results suggest that the peptides encoded by the *RAPTOR1B* and *MASP1* conserved non-AUG uORFs exert repressive effects on the expression of the mORF-encoded proteins. Consistent with these results, the *K*_a_/*K*_s_ analysis suggested that these two uORF sequences were conserved at the amino acid level rather than at the nucleotide level (Table 1).

**Figure 5.**
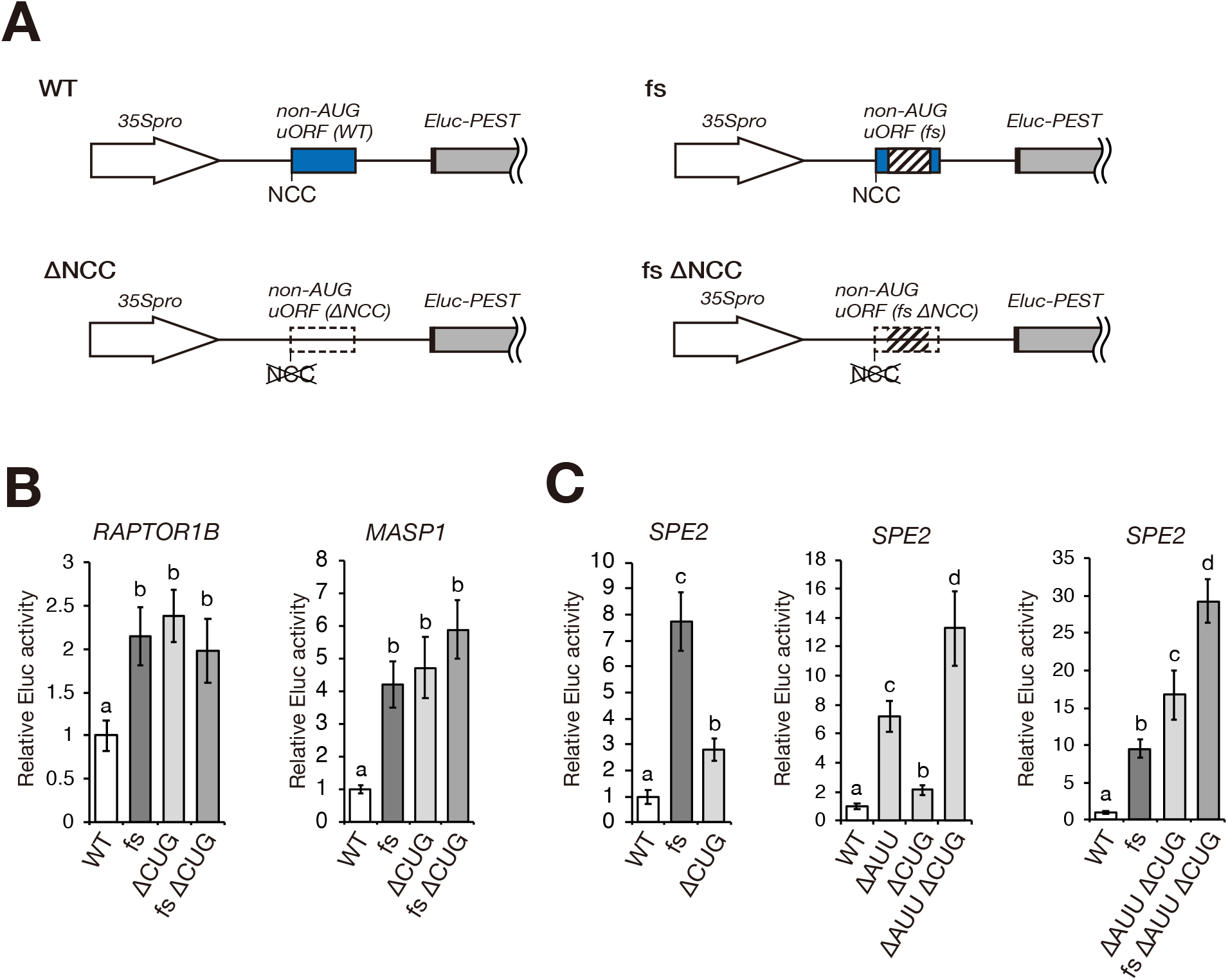
Sequence-dependent effects of the conserved non-AUG uORFs on mORF expression. (A) Schematic representation of the wild-type (WT) and mutant reporter constructs. The blue boxes in the WT and frameshift mutant (fs) constructs represent the conserved non-AUG uORFs. The open boxes with a dotted line in the ΔNCC and fs ΔNCC constructs depict the NCC-mutant non-AUG uORFs. The hatched boxes in the fs and fs ΔNCC construct show the frame-shifted region. See Supplementary Figure S3 for the exact position and length of the frame-shifted region of each conserved non-AUG uORF. (B) Transient expression assay. Each reporter plasmid containing the WT or mutant version of the 5′-UTR of the indicated gene was co-transfected into MM2d protoplasts with the *35S::Rluc* internal control plasmid. Eluc activity was normalized to Rluc activity. Data represent the mean ± SD of the normalized Eluc activities of at least three biological replicates, relative to that of the corresponding WT reporter construct. *P* values were calculated by Student’s *t*-test and adjusted by Bonferroni’s correction for multiple comparisons. Columns with different letters are significantly different at *P* < 0.05.

For *SPE2*, unexpectedly, the effect of the frameshift mutation was stronger than that of the NCC mutation (Figure 5C), indicating that the effect of the frameshift mutation was, at least in part, independent of translation from the conserved CUG codon. One possible explanation for this result is that translation of the *SPE2* non-AUG uORF initiates from another codon in addition to the conserved CUG codon. To test this possibility, we mutated the AUU codon located 9 nt upstream of the conserved CUG codon because this AUU codon was the only in-frame NCC located upstream of the conserved region (Supplementary Figure S3J). Although the AUU codon is not evolutionarily conserved, mutating the AUU codon more greatly increased reporter gene expression than the CUG mutation, and the AUU and CUG mutations showed an additive effect (Figure 5C). These results suggest that translation of the *SPE2* non-AUG uORF initiates from the AUU codon in addition to the conserved CUG codon. Consistent with this notion, ribosome profiling data show peaks of ribosome footprints around the AUU and CUG codons (Supplementary Figure S4). The AUU and CUG mutations and the frameshift mutation also exhibited an additive effect (Figure 5C), indicating that the effect of the frameshift mutation was independent of the translation from the AUU and CUG codons. This result does not support the idea that the amino acid sequence encoded by the *SPE2* non-AUG uORF is responsible for the repressive effect. Together with the high *K*_a_/*K*_s_ ratio of the *SPE2* non-AUG uORF (Table 1), these results suggest that translation from the AUU and CUG codons of the *SPE2* non-AUG uORF exerts a repressive effect on mORF expression but the repressive effect does not depend on the amino acid sequence of the *SPE2* non-AUG uORF.

### The *MPK22* non-AUG uORF alleviates the inhibitory effect of the downstream AUG CPuORF

In contrast to the *RAPTOR1B*, *MASP1* and *SPE2* uORFs, the *MPK22* non-AUG uORF was found to have a positive effect on mORF expression. This was unexpected because uORFs typically exert repressive effects on mORF translation. One possible mechanism to account for this is that, in *MPK22* mRNA, there is an inhibitory uORF downstream of the non-AUG uORF and the presence of the non-AUG uORF allows ribosomes to bypass the downstream inhibitory uORF, thereby promoting translation initiation at the mORF start codon. As mentioned in the Introduction, this role of uORFs in alleviating the effect of a downstream inhibitory effect has been observed in the AUG-initiated uORFs of the yeast *GCN4* gene and mammalian *ATF4* and *ATF5* genes (8–13). In the *MPK22* 5′-UTR, there is an AUG CPuORF downstream of the conserved non-AUG uORF (52), although the effect of the AUG CPuORF on mORF expression is not known. If the *MPK22* AUG CPuORF has an inhibitory effect and the conserved non-AUG uORF enhances mORF expression by promoting ribosome bypass of the AUG CPuORF, the effect of eliminating the conserved non-AUG codon should depend on the AUG CPuORF. To test this possibility, we mutated the start codon of the AUG CPuORF in the *MPK22* reporter construct and examined the effect of eliminating the conserved non-AUG uORF on reporter gene expression in the absence of the AUG CPuORF. As shown in Figure 6, the removal of the AUG CPuORF increased reporter gene expression by 10-fold, indicating that the *MPK22* AUG CPuORF has a strong inhibitory effect on mORF expression. In the absence of the AUG CPuORF, the removal of the conserved non-AUG uORF did not significantly affect reporter gene expression (Figure 6), indicating that the effect of the conserved non-AUG uORF on the expression of the *MPK22* mORF depends on the presence of the AUG CPuORF. This result suggests that, in *MPK22* mRNA, translation of the conserved non-AUG uORF directs ribosome bypass of the inhibitory AUG CPuORF, thereby promoting translation initiation at the mORF start codon.

**Figure 6.**
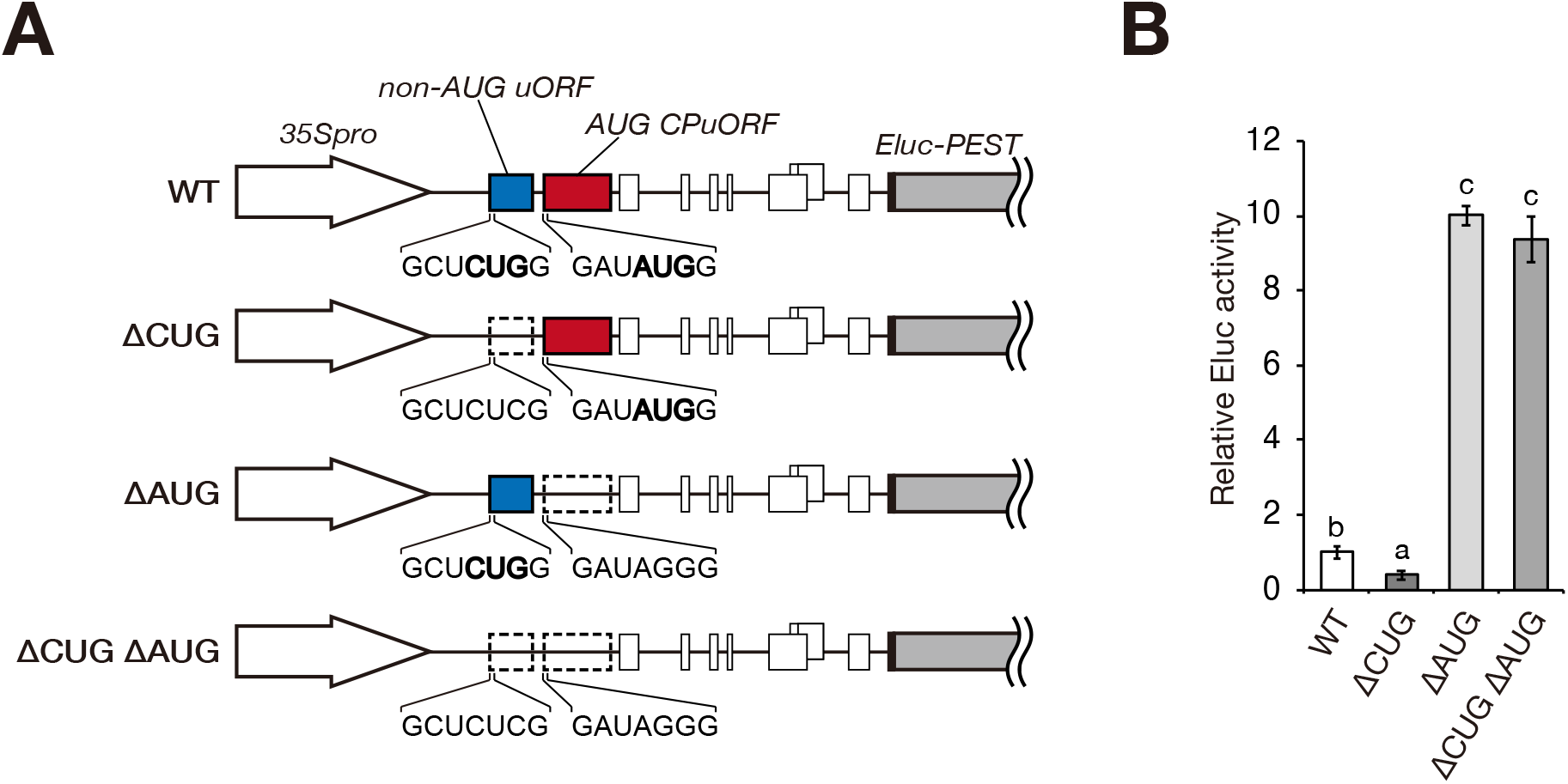
Effects of the *MPK22* conserved non-AUG and AUG uORFs on mORF expression. (A) Schematic representation of the wild-type (WT) and mutant reporter constructs. The WT construct contains the *MPK22* 5′-UTR between the 35S promoter (*35Spro*) and the *Eluc-PEST* coding sequence. The blue boxes represent the conserved CUG-initiated uORF. The red boxes show the AUG-initiated CPuORF. White boxes depict the other AUG-initiated uORFs. The CUG start codon of the non-AUG uORF, the AUG start codon of the AUG CPuORF, and their surrounding sequences are indicated. The start codons are shown in bold. The CUG start codon of the non-AUG uORF was mutated in the ΔCUG mutant. The start codon of the AUG CPuORF was mutated in the ΔAUG mutant. Both of these start codons were mutated in the ΔAUG mutant. The open boxes with a dotted line depict the mutant uORFs. (B) Transient expression assay. Each reporter plasmid containing the WT or mutant version of the *MPK22* 5′-UTR was co-transfected into MM2d protoplasts with the *35S::Rluc* internal control plasmid. Eluc activity was normalized to Rluc activity. Data represent the mean ± SD of the normalized Eluc activities of three biological replicates, relative to that of the WT reporter construct. *P* values were calculated by Student’s *t*-test and adjusted by Bonferroni’s correction for multiple comparisons. Columns with different letters are significantly different at *P* < 0.05.

## DISCUSSION

Interest in non-AUG uORFs has increased in recent years because of the identification of many translation initiation events at non-AUG codons in the 5′-UTRs of eukaryotic mRNAs by ribosome profiling studies. However, the physiological significance of the non-AUG translation initiation events in the 5′-UTRs is poorly understood, except for a few characterized non-AUG uORFs. In the present study, we searched for regulatory non-AUG uORFs with evolutionarily conserved sequences in Arabidopsis by combining bioinformatics and experimental approaches to identify physiologically important non-AUG uORFs. Bioinformatics analysis of the Arabidopsis genome identified 14 non-AUG uORFs with conserved NCCs and sequences, including four known conserved non-AUG uORFs. Transient expression assays identified four novel regulatory non-AUG uORFs and revealed that three of them exerted repressive effects on mORF expression, whereas one exerted a promotive effect. Two of the inhibitory non-AUG uORFs encode conserved amino acid sequence (35), and our assays showed that their inhibitory effects were sequence dependent. In addition to these, we identified three novel non-AUG CPuORFs (the *HEMA1*, *MYB7*, and *FH11* CPuORFs). However, our search did not identify 12 of the 14 non-AUG CPuORFs recently reported by van der Horst *et al*. (35). This is due to the differences in criteria for the selection of non-AUG CPuORFs (Supplementary Table S4). We discarded uORFs with an in-frame ATG codon upstream of the 5′-most NCC and uORFs with uORF-mORF fusion ratios equal to or greater than 0.3 in steps 1 and 3, respectively. Also, we discarded uORFs conserved only among less than five orders and uORFs without conserved NCCs in steps 5 and 6, respectively. Although we cannot rule out the possibility that some of these discarded uORFs have regulatory roles, we excluded these from candidates subjected to transient expression assays to focus on conserved non-AUG uORFs more likely to have a regulatory role.

Our transient expression assays showed that the conserved CUG codons are used as translation start codons for all of the four identified regulatory non-AUG uORFs, although translation of the *SPE2* uORF initiates from the unconserved AUU codon in addition to the conserved CUG codon. This is in agreement with previous observations that translation initiation at CUG is most efficient among NCCs (27, 28, 30). The conserved NCCs in three of the four regulatory non-AUG uORFs are in optimal initiation contexts, with a purine at ‒3 and a G at +4. This is also consistent with a previous report that translation initiation at NCCs is highly dependent on their sequence contexts, in particular the nucleotides at positions ‒3 and +4 (30). In contrast to these three non-AUG uORFs, the AUU and CUG start codons of the *SPE2* non-AUG uORF are in non-optimal initiation contexts. The *SPE2* 5′-UTR contains a hairpin structure, which is widely conserved across land plants (53), 19 nt downstream of the CUG start codon (Supplementary Figure S4). There are some known examples in which a hairpin RNA structure promotes translation initiation at an NCC by causing PICs to stall near the NCC. An earlier study using a rabbit reticulocyte lysate cell-free translation system found that translation initiation at a GUG codon is promoted by a hairpin structure beginning 15 nt downstream of the GUG codon (54). More recently, translation initiation from an AUU codon of the mORF in human *PTEN* mRNA was shown to require a hairpin structure that started 18 nt downstream of the AUU codon (55). Also, in human *POLG* mRNA, translation of the mORF is efficiently initiated at a CUG codon located 14 nt upstream of a hairpin structure (56). The region including the hairpin structure in the *SPE2* 5′-UTR has been reported to be important for translational repression of the mORF (53). Therefore, it is likely that the hairpin structure helps PICs recognize the AUU and CUG start codons in the non-optimal sequence contexts, thereby exerting the repressive effect on mORF translation. The *SPE2* mORF codes for Arg decarboxylase, a key enzyme in polyamine biosynthesis. Polyamines are known to affect RNA secondary structures (57). Therefore, if translation initiation at the conserved CUG codon of the *SPE2* non-AUG uORF depends on the hairpin structure and cellular polyamine concentration affects the hairpin structure, the *SPE2* non-AUG uORF could be involved in feedback regulation of polyamine biosynthesis.

Of the four regulatory non-AUG uORFs identified in this study, the *RAPTO1B* and *MASP1* non-AUG uORFs exhibited sequence-dependent repressive effects on mORF expression. In contrast to the significant effects of the NCC mutations introduced into the WT version of these two uORFs, mutating the conserved NCCs of the frameshift mutant version of these uORFs had no significant effect (Figure 5B). This suggests that the translation initiation efficiencies at the conserved CUG codons of these uORFs are not high enough to exert significant repressive effects simply by capturing ribosomes, but that the repressive effects are enhanced by the uORF-encoded peptide sequences, most likely via ribosome stalling. In eukaryotes, more than 30 AUG uORFs have been reported to exert repressive effects on mORF expression in an amino acid sequence-dependent manner (14, 16, 38, 41, 58–60). In contrast, only two sequence-dependent regulatory non-AUG uORFs have been previously reported (21, 22). Both of these two non-AUG uORFs play a role in feedback regulation of metabolite biosynthesis. On the other hand, *MASP1* and *RAPTO1B* code for proteins involved in stress responses. MASP1 expression is upregulated in response to drought stress, and this upregulation occurs at the protein level rather than at the mRNA level (61). Therefore, the sequence-dependent regulatory non-AUG uORF could be involved in the drought stress-responsive translational regulation of *MASP1*. It remains to be elucidated why CUG, and not AUG, is used as the start codons of the *MASP1* and *RAPTO1B* uORFs. One possibility is that the low translation initiation efficiency of CUG compared to AUG is favorable for repressing mORF expression only under conditions where ribosome stalling occurs, as seen in the translational regulation of the *N. crassa arg2* gene, which involves Arg-responsive ribosome stalling mediated by a uORF starting with an AUG codon in a poor initiation context (15). Another possibility is that the efficiencies of translation initiation at the conserved CUG start codons of the *MASP1* and *RAPTO1B* non-AUG uORFs change depending on environmental or cellular conditions, thereby controlling mORF expression in a condition-dependent manner. In mammals, two examples of stress-induced changes in translation initiation efficiencies of non-AUG uORFs have been reported. In the *EPRS* mRNA encoding a glutamyl-proryl-tRNA synthetase, the translation initiation efficiencies of two NCC-initiated uORFs are reduced under stress conditions where the α subunit of eIF2 (eIF2α is phosphorylated, resulting in enhanced translation of the mORF (24). In another example, translation of two non-AUG uORFs in *BiP* mRNA is promoted under endoplasmic reticulum stress conditions, depending on the monomeric eukaryotic initiation factor eIF2A (23). eIF2A is required for translation initiation with Leu-tRNA (62), and its expression level is increased in response to various stresses (23, 63, 64). Although such an alternative eukaryotic initiation factor that specifically promotes translation initiation at NCCs has not been reported in plants, it is likely that plants have similar mechanisms to regulate translation initiation at NCCs in response to stresses. This is because the stress-induced phosphorylation of eIF2α is conserved in plants (65–68), and the Arabidopsis genome contains a gene encoding an amino acid sequence similar to that of mammalian eIF2A although their functional similarities have not yet been experimentally validated.

In contrast to the other regulatory non-AUG uORFs identified in this study, the *MPK22* non-AUG uORF promotes mORF expression. Some non-AUG uORFs with positive effects on mORF translation have been previously reported (23, 36). In addition, genome-wide analysis of yeast ribosome profiling data revealed that genes with NCC-initiated uORFs had significantly higher translation efficiencies than genes without uORFs, whereas genes with AUG-initiated uORFs had significantly lower translation efficiencies than genes without uORFs (36). However, the molecular mechanisms by which these non-AUG uORFs promote mORF translation remain to be elucidated. Our data revealed that the effect of the *MPK22* non-AUG uORF depended on the downstream inhibitory AUG CPuORF (Figure 6), suggesting that the non-AUG uORF alleviates the repressive effect of the AUG CPuORF by allowing ribosomes to bypass the AUG CPuORF. In eukaryotes, some AUG uORFs are known to derepress mORF expression under specific conditions by alleviating the repressive effect of a downstream uORF (8–13), whereas no non-AUG uORF that derepresses mORF expression in this way has been previously reported. Our results demonstrated that a non-AUG uORF is able to exert a significant derepressive effect on mORF expression by alleviating the effect of a downstream inhibitory uORF if the downstream uORF has a strong repressive effect, although the derepressive effect of the non-AUG uORF would be weaker than that of an AUG uORF with a high translation initiation efficiency. This unveiled a mechanism by which a non-AUG uORF can promote mORF expression, although this mechanism alone cannot explain the positive effects of all other non-AUG uORFs on mORF expression. It remains to be elucidated why a non-AUG uORF, and not an AUG uORF, is used in *MPK22* to alleviate the repressive effect of the AUG CPuORF. One possibility is that the non-AUG uORF is used to fine-tune expression from the *MPK22* mORF at an appropriate level. If this is the case, alleviation by an AUG uORF may cause the MPK22 expression level to be too high, and therefore, the non-AUG uORF may be preferable to moderately alleviate the repressive effect of the downstream AUG CPuORF. Another possibility is that the non-AUG uORF is involved in condition-dependent regulation of MPK22 expression. For example, if certain conditions increase the translation initiation efficiency of the *MPK22* non-AUG uORF, the alleviation of the inhibitory effect of the AUG CPuORF by the non-AUG uORF would be enhanced under these conditions, thereby increasing the MPK22 expression level. The *MPK22* mORF codes for a MAP kinase. Although the conditions under which this kinase is activated are not yet known, other plant MAP kinases are known to be activated in response to biotic or abiotic stresses or phytohormones (69). Therefore, it is possible that MPK22 expression as well as its activity are activated under specific conditions, and that the non-AUG uORF and the AUG CPuORF are involved in the condition-dependent regulation of MPK22 expression.

## Conclusion

This study identified four sequence-dependent and -independent regulatory non-AUG uORFs in Arabidopsis. Through the analysis of one of the sequence-independent regulatory non-AUG uORFs (the *MPK22* non-AUG uORF), we discovered a new type of regulatory non-AUG uORF, which promotes mORF expression by alleviating the inhibitory effect of a downstream uORF. In addition, while only two sequence-dependent regulatory non-AUG uORFs have been previously reported in eukaryotes, the current study identified two novel sequence-dependent regulatory non-AUG uORFs. These two types of mechanisms enable non-AUG-uORFs to exert significant effects on mORF expression. Although the physiological roles of the newly identified regulatory non-AUG uORFs remain to be elucidated, their evolutionarily conserved sequences imply that these non-AUG uORFs play important regulatory roles. Further physiological studies on translational regulation mediated by these non-AUG uORFs will provide a better understanding of the regulatory roles of non-AUG uORFs.

## SUPPLEMENTARY DATA

**Supplementary Table S1.** Plasmids used in this study and primers used for plasmid construction.

**Supplementary Table S2.** Primers used in this study.

**Supplementary Table S3.** Taxonomic range of sequence conservation of the conserved non-AUG uORFs identified in this study.

**Supplementary Table S4.** Identification of previously discovered non-AUG CPuORFs.

**Supplementary Figure S1.** Schematic representation of BLAST-based search for uORFs conserved between homologous genes.

**Supplementary Figure S2.** Alignments of the novel and known conserved non-AUG uORFs identified in this study.

**Supplementary Figure S3.** Nucleotide sequences of the 5′-UTRs used for the transient expression study and the deduced amino acid sequences of the conserved non-AUG uORFs.

**Supplementary Figure S4.** Ribosome footprint distribution along the *SPE2* mRNA.

## ACKNOWLEDGMENTS

We thank Ms. Kazuko Harada and Ms. Sayo Aono for general assistance. We used the DNA sequencing facility of the Graduate School of Agriculture, Hokkaido University.

