## Supplementary data for "Genome-wide identification of Arabidopsis non-AUG-initiated upstream ORFs with evolutionarily conserved regulatory sequences that control protein expression levels"

Supplementary Table S1. Plasmids used in this study and primers used for plasmid construction

| Plasmid | Construct | Primer | |
|---|---|---|---|
| | | Forward | Reverse |
| pKM5 | *35S::Rluc* | – | – |
| pKM56 | *35S::Eluc-PEST* | – | – |
| pYH1 | *35S::KAN2(WT):Eluc-PEST* | KAN2 F | KAN2 R |
| pYH2 | *35S::KAN2(ΔGUG):Eluc-PEST* | KAN2 d1 F | KAN2 d1 R |
| pYH3 | *35S::TTL1(WT):Eluc-PEST* | nTTL1 F | TTL1 R |
| pYH4 | *35S::TTL1(ΔAUU):Eluc-PEST* | TTL1 d F | TTL1 d R |
| pYH5 | *35S::HEMA1(WT):Eluc-PEST* | HEMA1 F | HEMA1 R |
| pYH6 | *35S::HEMA1(ΔGUG):Eluc-PEST* | HEMA1 dGTG F | HEMA1 dGTG R |
| pYH7 | *35S::MYB7(WT):Eluc-PEST* | MYB7 F | MYB7 R |
| pYH8 | *35S::MYB7(ΔGUG):Eluc-PEST* | MYB7 d F | MYB7 d R |
| pYH9 | *35S::SPEAR1(WT):Eluc-PEST* | SPEAR1 F | SPEAR1 R |
| pYH10 | *35S::SPEAR1(ΔAUC):Eluc-PEST* | SPEAR1 nd F | SPEAR1 R |
| pYH11 | *35S::FH11(WT):Eluc-PEST* | FH11 F | FH11 R |
| pYH12 | *35S::FH11(ΔUUG):Eluc-PEST* | FH11 d F | FH11 d R |
| pYH13 | *35S::RAPTOR1B(WT):Eluc-PEST* | RAPTOR F | RAPTOR R |
| pYH14 | *35S::RAPTOR1B(ΔCUG):Eluc-PEST* | RAPTOR CCC F | RAPTOR CCC R |
| pYH15 | *35S::MASP1(WT):Eluc-PEST* | MASP1 Xba1 SLiCE F | MASP1 Sal1 SLiCE R |
| pYH16 | *35S::MASP1(ΔCUG):Eluc-PEST* | nMASP1 CTC F | nMASP1 CTC R |
| pYH17 | *35S::MPK22(WT):Eluc-PEST* | MPK22 F | MPK22 R |
| pYH18 | *35S::MPK22(ΔCUG):Eluc-PEST* | MPK22 d2 F | MPK22 d2 R |
| pYH19 | *35S::SPE2(WT):Eluc-PEST* | SPE2 F | SPE2 R |
| pYH20 | *35S::SPE2(ΔCUG):Eluc-PEST* | SPE2 d F | SPE2 d R |
| pYH21 | *35S::APD6(WT):Eluc-PEST* | APD6 F | APD6 R |
| pYH22 | *35S::APD6(ΔGUG):Eluc-PEST* | APD6 d1 F | APD6 R |
| pYH23 | *35S::AHK2(WT):Eluc-PEST* | AHK2 F | AHK2 R |
| pYH24 | *35S::AHK2(ΔAUU):Eluc-PEST* | AHK2 d2 F | AHK2 d2 R |
| pYH25 | *35S::TcMYB7(WT):Eluc-PEST* | – | – |
| pYH26 | *35S::TcMYB7(ΔCUG):Eluc-PEST* | cacao MYB7 dCUG F | cacao MYB7 dCUG R |
| pYH27 | *35S::AiFH11(WT):Eluc-PEST* | – | – |
| pYH28 | *35S::AiFH11(ΔCUG):Eluc-PEST* | Archis FH11 dCUG F | Archis FH11 dCUG R |
| pYH29 | *35S::TcAPD6(WT):Eluc-PEST* | – | – |
| pYH30 | *35S::TcAPD6(ΔUUG):Eluc-PEST* | Cacao APD6 dUUG F | Cacao APD6 dUUG R |
| pYH31 | *35S::TcAPD6(ΔCUG):Eluc-PEST* | Cacao APD6 dCUG F | Cacao APD6 dCUG R |
| pYH32 | *35S::TcAPD6(ΔAUU):Eluc-PEST* | Cacao APD6 dAUU F | Cacao APD6 dAUU R |
| pYH33 | *35S::HEMA1(fs):Eluc-PEST* | HEMA1 nnfs F | HEMA1 nnfs R |
| pYH34 | *35S::HEMA1(fs ΔGUG):Eluc-PEST* | HEMA1 dGTG F | HEMA1 dGTG R |
| pYH35 | *35S::RAPTOR1B(fs):Eluc-PEST* | RAPTOR fs F | RAPTOR fs R |
| pYH36 | *35S::RAPTOR1B(fs ΔCUG):Eluc-PEST* | RAPTOR CCC F | RAPTOR CCC R |
| pYH37 | *35S::MASP1(fs):Eluc-PEST* | MASP1 fs2 F | MASP1 fs2 R |
| pYH38 | *35S::MASP1(fs ΔCUG):Eluc-PEST* | nMASP1 CTC F | nMASP1 CTC R |
| pYH39 | *35S::SPE2(fs):Eluc-PEST* | SPE2 nfs F | SPE2 nfs R |
| pYH40 | *35S::SPE2(fs ΔCUG):Eluc-PEST* | SPE2 d F | SPE2 d R |
| pYH41 | *35S::MPK22(ΔAUG):Eluc-PEST* | MPK22 dCPuORF25 F | MPK22 dCPuORF25 R |
| pYH42 | *35S::MPK22(ΔCUG ΔAUG):Eluc-PEST* | MPK22 dCPuORF25 F | MPK22 dCPuORF25 R |

**Supplementary Table S2. Primers used in this study**

| Name | Primer sequence |
|---|---|
| KAN2 F | AGAACACGGGGGACTCTAGAAGCCCCCCATGTCCACAC |
| KAN2 R | TCTTCTCTCTCTCCGTCGACTCCATGAAAGAGACCTTTAACTTC |
| KAN2 d1 F | CACACACTCAAGTATCATTTTCTCA |
| KAN2 d1 R | TGAGAAAATGATACTTGAGTGTGTG |
| nTTL1 F | AGAACACGGGGGACTCTAGAGGTTCTCCATCTTCACTCCTC |
| TTL1 R | TCTTCTCTCTCTCCGTCGACGGCATTTTGAGTGTTGTGGTGA |
| TTL1 d F | TCACATTCGTTCTTTGCTTCAAC |
| TTL1 d R | AAGCAAAGAATGAACGTGAGTGC |
| HEMA1 F | AGAACACGGGGGACTCTAGATGTTGCCGTGTAAGAACAAATGCC |
| HEMA1 R | TCTTCTCTCTCTCCGTCGACACCGCCATTGAAACCCAAAATCTC |
| HEMA1 dGTG F | GCAAAAACGAGTGTCGATAAAACGTG |
| HEMA1 dGTG R | CACGTTTTATCGACACTCGTTTTTGC |
| MYB7 F | AGAACACGGGGGACTCTAGAATAATGGTTATGAGATCTAAAACATCA |
| MYB7 R | TCTTCTCTCTCTCCGTCGACCCCATGACTCTCTTCTTCTGA |
| MYB7 d F | CCTTCAAACGTCTCCTACCAAC |
| MYB7 d R | GGTAGGAGACGTTTGAAGGA |
| SPEAR1 F | AGAACACGGGGGACTCTAGAGGCGTATCTATTGAAATCTGGAA |
| SPEAR1 R | TCTTCTCTCTCTCCGTCGACCCCATGTTTCTTTGATCGGC |
| SPEAR1 nd F | AGAACACGGGGACTCTAGAGGCGTATCTATTGAACTCTGGA |
| FH11 F | AGAACACGGGGGACTCTAGAGCCAACAATAAGAGGACCCA |
| FH11 R | TCTTCTCTCTCTCCGTCGACACCATATCTTTGTTCTCTTTTTAGATC |
| FH11 d F | CCAGTACAACTATAGTTACAGAGTTCA |
| FH11 d R | TGAACTCTGTAACTATAGTTGTACTGG |
| RAPTOR F | GAGAGAACACGGGGGACTCTAGAGAAGTGGAAAGAAAACGCAA |
| RAPTOR R | CGTTCTTCTCTCTCTCCGTCGACGCCATCCAAATCGGAGAAC |
| RAPTOR CCC F | GGGTTTTTCGAGCCCGATTCTCTGCTC |
| RAPTOR CCC R | GAGCAGAGAATCGGGCTCGAAAAACCC |
| MASP1 Xba1 SLiCE F | AGAACACGGGGACTCTAGAGCGTTGCTTTGCCTCAA |
| MASP1 Sal1 SLiCE R | CGTTCTTCTCTCTCTCCGTCGACAACCTGACCATTGTGTCAAAC |
| nMASP1 CTC F | TCTTCACTCTCGCCGCTTCTGC |
| nMASP1 CTC R | AGAAGCGGCGAGAGTGAAGAAC |
| MPK22 F | AGAACACGGGGGACTCTAGAACGAGAGAACCGCACATTGA |
| MPK22 R | TCTTCTCTCTCTCCGTCGACTCCATCTTTTTTTTTCTGGTTAA |
| MPK22 d2 F | GAGAAGCTCTCGATTCCGTC |
| MPK22 d2 R | GACGGAATCGAGAGCTTCTC |
| SPE2 F | AGAACACGGGGGACTCTAGAACGCCTTCTCTTCTTCTTCTTC |
| SPE2 R | TCTTCTCTCTCTCCGTCGACGGCATCTTTATCTTCACCCTC |
| SPE2 d F | GATTCATCTTCTCTCTTCTCCTGA |
| SPE2 d R | TCAGGAGAAGAGAGAAGATGAATC |
| APD6 F | AGAACACGGGGGACTCTAGATGGAGCCCGAAAGCTGGG |
| APD6 R | TCTTCTCTCTCTCCGTCGACAGCATCCCTCATCTCATCC |
| APD6 dintron F | GCTTCCAACATCTTGCAAGTTGTCTTTGAGTTCATTCAGGTC |
| APD6 dintron R | CTTGCAAGATGTTGGAAGC |
| APD6 d1 F | AGAACACGGGGACTCTAGATGGAGCCCGAAAGCTGGGACCTTTGATGTCCTGGCTCC |
| AHK2 F | AGAACACGGGGGACTCTAGAATAATTTCTCTAACAAATGCTTCTTTTA |
| AHK2 R | TCTTCTCTCTCTCCGTCGACGACATTTCGACTCCTAATCTCA |
| AHK2 d2 F | ACTCTGATCGTTGAGTCCAGTTG |
| AHK2 d2 R | CCAACTGGACTCAACGATCAGA |

**Supplementary Table S2. Primers used in this study (continued)**

| Name | Primer sequence |
|---|---|
| cacao MYB7 dCUG F | CCTCAAACCTCTCTCACCAA |
| cacao MYB7 dCUG R | TTGGTGAGAGAGGTTTGAGG |
| Archis FH11 dCUG F | AACCACAACTCCCAGGTTCA |
| Archis FH11 dCUG R | TGAACCTGGGAGTTGTGGTT |
| cacaoAPD6 dUUG F | CCTTTGCTTTACTGGCTTTAA |
| cacaoAPD6 dUUG R | TTAAAGCCAGTAAAGCAAAGG |
| Cacao APD6 dUUG F | CTTTGCTTTCCTGGCTTTA |
| Cacao APD6 dUUG R | TAAAGCCAGGAAAGCAAAG |
| Cacao APD6 dCUG F | TGCTTTGCTCGCTTTAAAC |
| Cacao APD6 dCUG R | GTTTAAAGCGAGCAAAGCA |
| Cacao APD6 dAUU F | TGCTGAGAGTTGGAGGGA |
| Cacao APD6 dAUU R | TCCCTCCAACTCTCAGCA |
| HEMA1 nnfs F | CTCCACCCGCCAATCAAATTCCATCAAATCTCAATGC |
| HEMA1 nnfs R | TTTGATTGGCGGGTGGAGAACACGTTTTTATCCAC |
| RAPTOR fs F | CGCTGGTGCAGCGGCTTTGTGATTGATC |
| RAPTOR fs R | AAAGCCGCTGCACCAGCGCGCGCAGGAGC |
| MASP1 fs2 F | TGTTTCTGAGGGTCTTACCCTTGATTACGGCTTAACATTTAC |
| MASP1 fs2 R | GGGTAAGACCCTCAGAAACACGACGGAGCCTCCAATGATAGA |
| SPE2 nfs F | CCGGCCTCGGCGGTTTTTTAAACCCCTACCTTTACCAAATTCTGG |
| SPE2 nfs R | AAAAACCGCCGAGGCCGGAGCCTCGGCTACCCCCAGGAGAAGAC |
| MPK22 dCPuORF25 F | TCATCTTCTGAGATAGGGAACAAGT |
| MPK22 dCPuORF25 R | ACTTGTTCCCTATCTCAGAAGATGA |
| Xba1 SLiCE F | AGAACACGGGGACTCTAGA |
| Sal1 SLiCE R | TCTTCTCTCTCTCCGTCGAC |
| Spe1 SLiCE F | AGAACACGGGGACTCTAGT |

**Supplementary Table S3. Taxonomic range of sequence conservation of the conserved non-AUG uORFs identified in this study.**

| Gene ID (AGI code) | Gene symbol | Taxonomic category ** | | | | | | | | | | | | |
|---|---|---|---|---|---|---|---|---|---|---|---|---|---|---|
| | | Lamiids | Asterids* | Malvids | Fabids | Eudicots* | Commelinids | Monocots* | Angiospermae* | Gymnospermae | Polypodiopsida | Embryophyta* | Streptophyta* | Viridiplantae* |
| AT1G32240 | KAN2 | 2 | 4 | 3 | 6 | 4 | 0 | 0 | 0 | 0 | 0 | 0 | 0 | 0 |
| AT1G53300 | TTL1 | 2 | 1 | 1 | 3 | 2 | 0 | 1 | 0 | 0 | 0 | 0 | 0 | 0 |
| AT1G58290 | HEMA1 | 0 | 1 | 1 | 2 | 2 | 0 | 0 | 0 | 0 | 0 | 0 | 0 | 0 |
| AT2G16720 | MYB7 | 2 | 2 | 1 | 3 | 1 | 0 | 0 | 0 | 0 | 0 | 0 | 0 | 0 |
| AT2G20080 | SPEAR1 | 0 | 0 | 1 | 3 | 0 | 0 | 0 | 0 | 0 | 0 | 0 | 0 | 0 |
| AT3G05470 | FH11 | 1 | 0 | 2 | 3 | 2 | 0 | 0 | 0 | 0 | 0 | 0 | 0 | 0 |
| AT3G08850 | RAPTOR1B | 1 | 5 | 2 | 3 | 4 | 2 | 3 | 0 | 0 | 0 | 0 | 0 | 0 |
| AT4G03260 | MASP1 | 3 | 5 | 3 | 7 | 5 | 3 | 5 | 1 | 4 | 0 | 1 | 1 | 0 |
| AT4G19110 | MPK22 | 1 | 0 | 1 | 3 | 1 | 0 | 1 | 0 | 0 | 0 | 0 | 0 | 0 |
| AT4G26850 | VTC2 | 4 | 5 | 3 | 6 | 6 | 3 | 5 | 3 | 4 | 2 | 1 | 3 | 4 |
| AT4G34710 | SPE2 | 1 | 0 | 1 | 1 | 1 | 0 | 0 | 0 | 0 | 0 | 0 | 0 | 0 |
| AT4G38520 | APD6 | 0 | 0 | 1 | 5 | 2 | 0 | 1 | 0 | 0 | 0 | 0 | 0 | 0 |
| AT5G35750 | AHK2 | 2 | 0 | 1 | 1 | 2 | 0 | 0 | 0 | 0 | 0 | 0 | 0 | 0 |
| AT5G55120 | VTC5 | 4 | 5 | 3 | 6 | 6 | 3 | 5 | 3 | 4 | 2 | 1 | 3 | 4 |

** Numbers of orders from which uORF-mORF homologs were identified are indicated. The presence of the uORF-mORF homologs in each of the 13 taxonomic categories is indicated by highlighting cells in yellow.

* Orders included in lower taxonomic categories were excluded.

**Supplementary Table S4. Identification of previously discovered non-AUG CPuORFs.**

| Gene ID (AGI code) | Gene symbol | Reference | Identification in the steps in the present study | | | | | | |
|---|---|---|---|---|---|---|---|---|---|
| | | | Step 1 | Step 2 | Step 3 | Step 4 | Step 5 | Step 6 | Step 7 |
| AT1G01060 | LHY | 35 | ✓ | ✓ | ✓ | ✓ | NI | - | - |
| AT1G07640 | OBP2 | 35 | ✓ | ✓ | ✓ | ✓ | NI | - | - |
| AT1G66540 | AT1G66540 | 35 | ✓ | ✓ | NI | - | - | - | - |
| AT1G72820 | AT1G72820 | 35 | ✓ | NI | - | - | - | - | - |
| AT1G77840 | AT1G77840 | 35 | ✓ | ✓ | ✓ | ✓ | ✓ | ✓ | NI |
| AT2G23570 | MES19 | 35 | NI | - | - | - | - | - | - |
| AT2G29290 | AT2G29290 | 35 | NI | - | - | - | - | - | - |
| AT3G08730 | PK1 | 35 | ✓ | ✓ | ✓ | ✓ | ✓ | NI | - |
| AT3G08850 | RAPTOR1B | 35 | ✓ | ✓ | ✓ | ✓ | ✓ | ✓ | ✓ |
| AT3G23010 | RLP36 | 35 | ✓ | ✓ | NI | - | - | - | - |
| AT4G03260 | MASP1 | 35 | ✓ | ✓ | ✓ | ✓ | ✓ | ✓ | ✓ |
| AT4G26850 | VTC2 | 22 | ✓ | ✓ | ✓ | ✓ | ✓ | ✓ | ✓ |
| AT5G26140 | LOG9 | 35 | ✓ | ✓ | NI | - | - | - | - |
| AT5G55120 | VTC5 | 22 | ✓ | ✓ | ✓ | ✓ | ✓ | ✓ | ✓ |
| AT5G63190 | AT5G63190 | 35 | ✓ | ✓ | ✓ | ✓ | ✓ | NI | - |

NI, not identified.

### Supplementary Figure S1

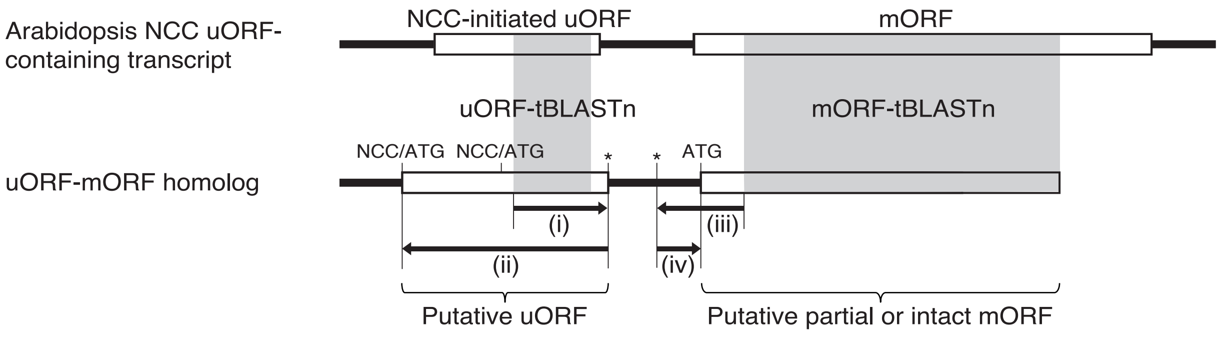


**Supplementary Figure S1.** Schematic representation of BLAST-based search for uORFs conserved between homologous genes. In step 4 of the genome-wide search for conserved NCC-initiated uORFs in this study, tBLASTn searches were conducted against a transcript sequence database that consists of assembled EST/TSA contigs, unclustered singleton EST/TSA sequences and RefSeq RNAs, using Arabidopsis NCC-initiated uORF sequences as queries (uORF-tBLASTn). The shaded regions in the open boxes show the tBLASTn-matching regions. Asterisks represent stop codons. (i) The downstream in-frame stop codon closest to the 5'-end of the matching region of each uORF-tBLASTn hit was selected. (ii) The 5'-most in-frame ATG codon or NCC other than AAG and AGG (NCC/ATG) located upstream of the stop codon was selected. The ORF beginning with the selected NCC/ATG and ending with the selected stop codon was extracted as a putative uORF. In step 5, the downstream sequences of putative uORFs in the transcript sequences were subjected to mORF-tBLASTn analysis. Transcript sequences matching the mORF of the original Arabidopsis uORF-containing transcript with an *E*-value less than $10^{-1}$ were extracted. (iii) For each of the mORF-tBLASTn hits, the upstream in-frame stop codon closest to the 5'-end of the matching region was selected. (iv) The 5'-most in-frame ATG codon located downstream of the selected stop codon was identified as the start codon of the putative partial or intact mORF. If the putative mORF overlaps with the putative uORF, the uORF-tBLASTn and mORF-tBLASTn hit is discarded as a uORF-mORF fusion type.

### Supplementary Figure S2

#### *AT1G32240 KAN2*

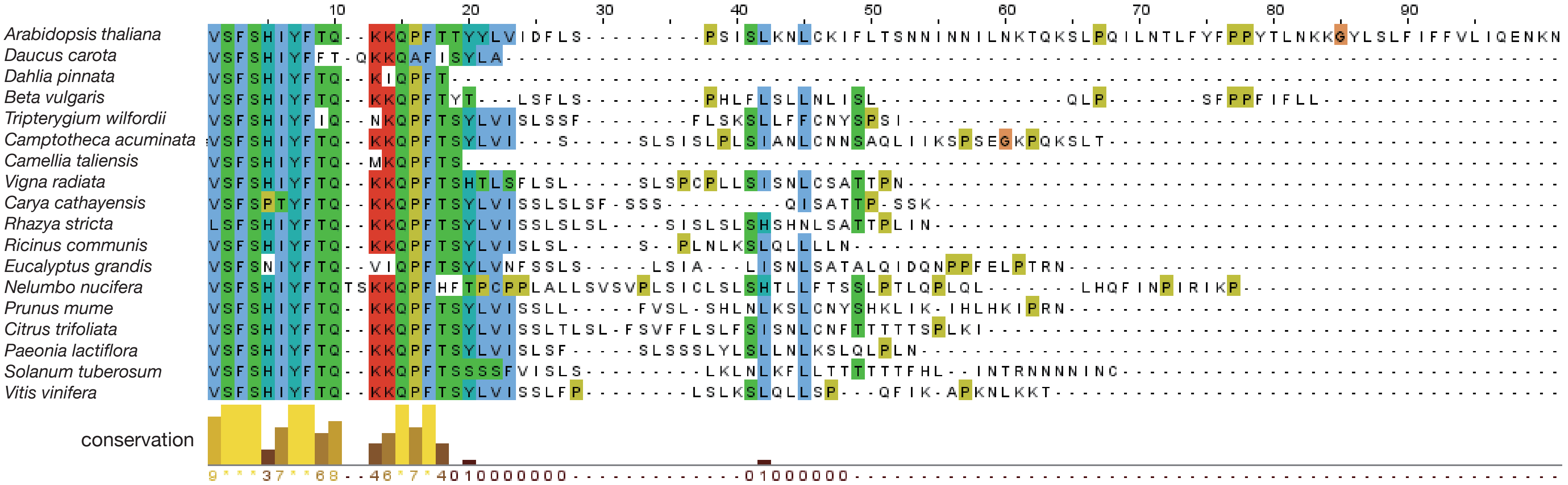


#### AT1G53300 *TTL1*

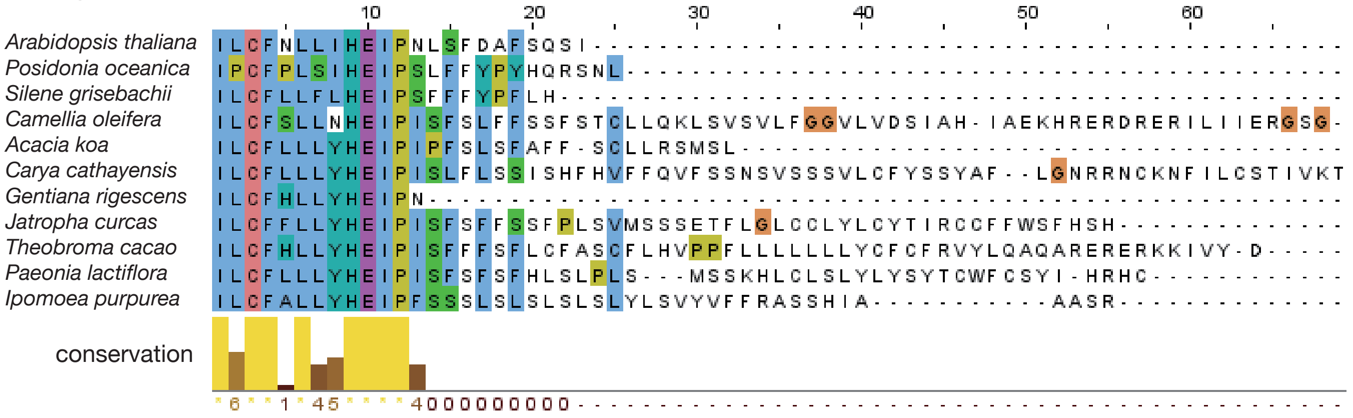

#### AT1G58290 *HEMA1*

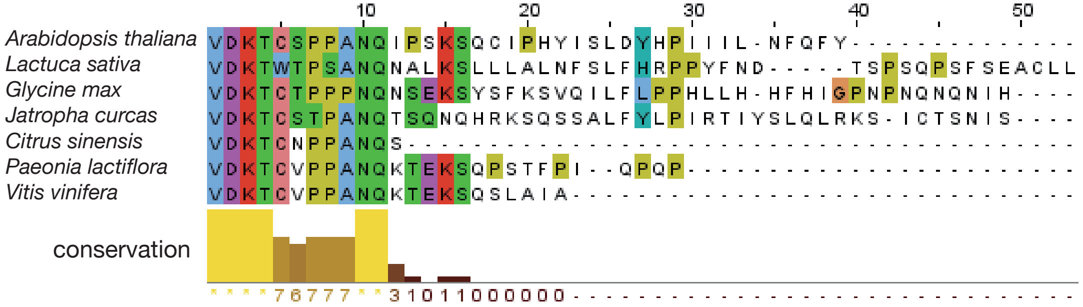


#### AT2G16720 *MYB7*

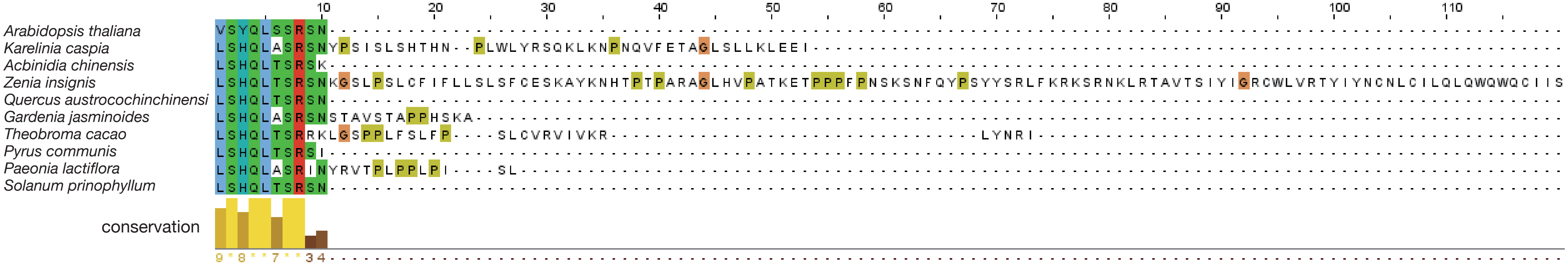


#### AT2G20080 *SPEAR1*

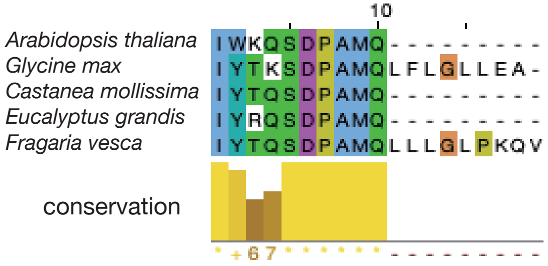

## AT3G05470 *FH11*

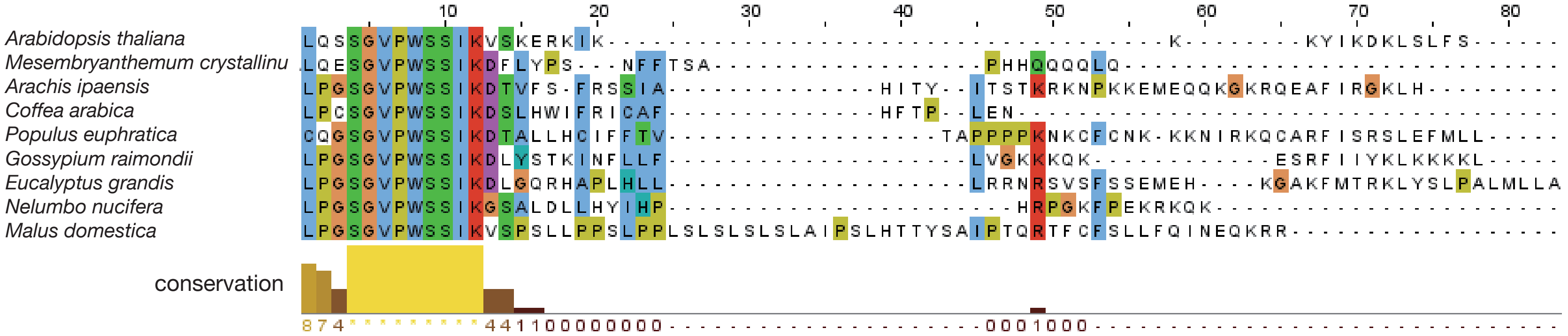


#### AT3G08850 *RAPTOR1B*

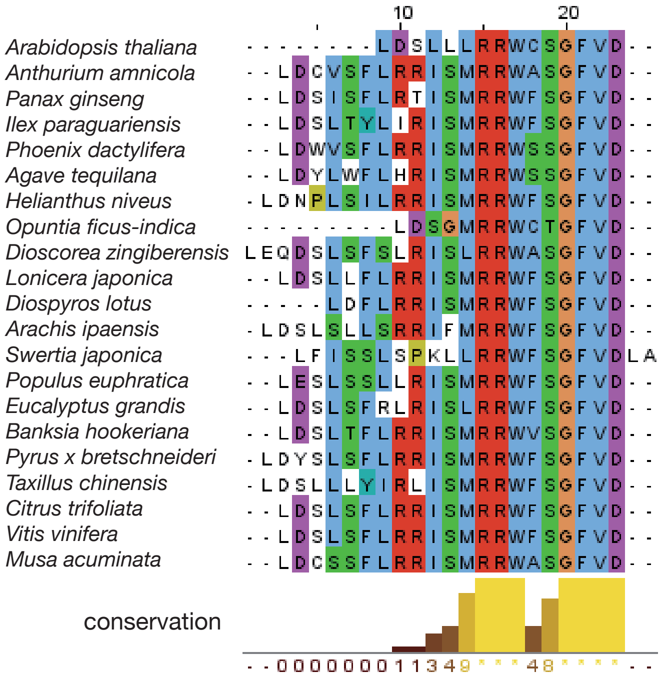

#### AT4G03260 *MASP1*

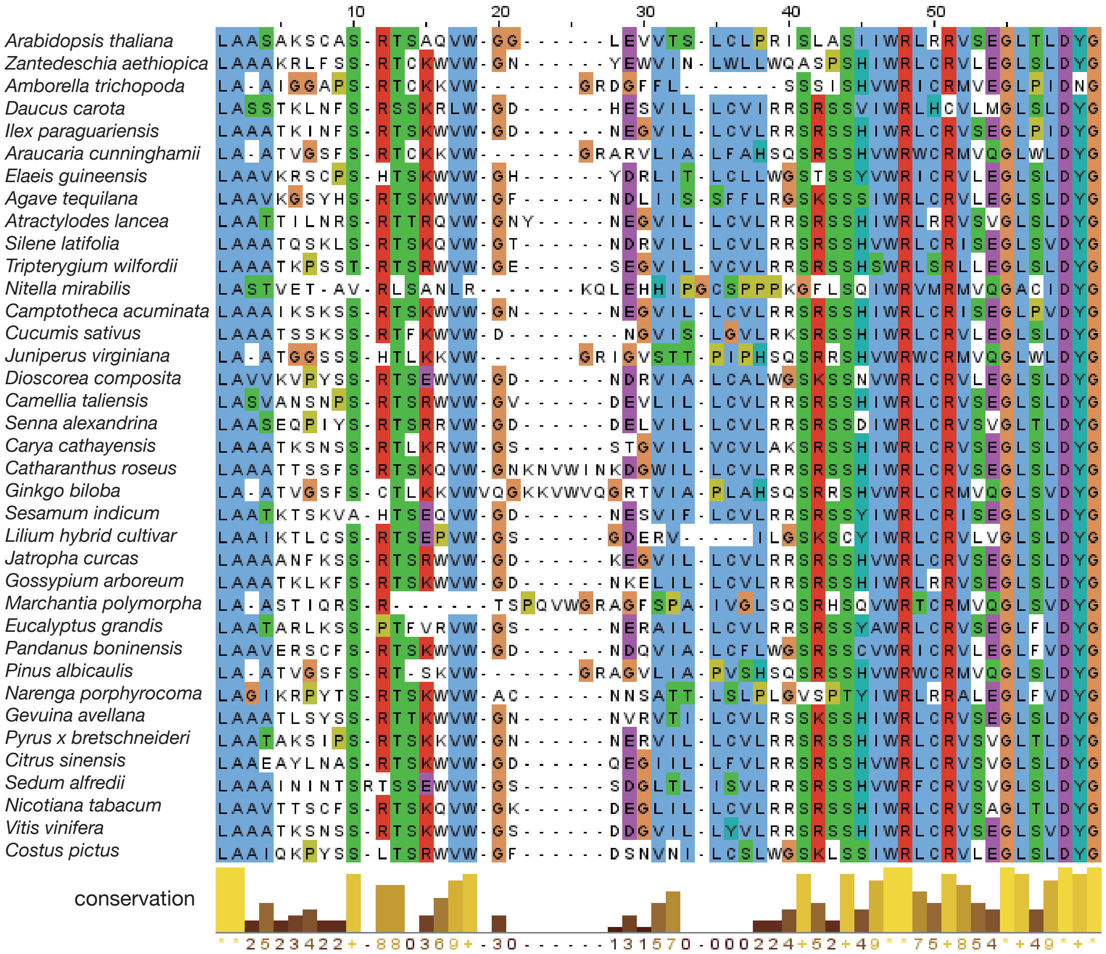


#### AT4G19110 *MPK22*

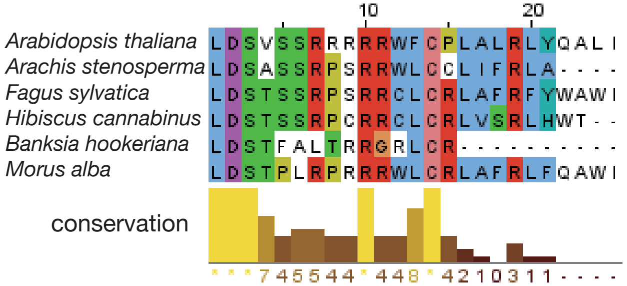

#### AT4G26850 *VTC2*

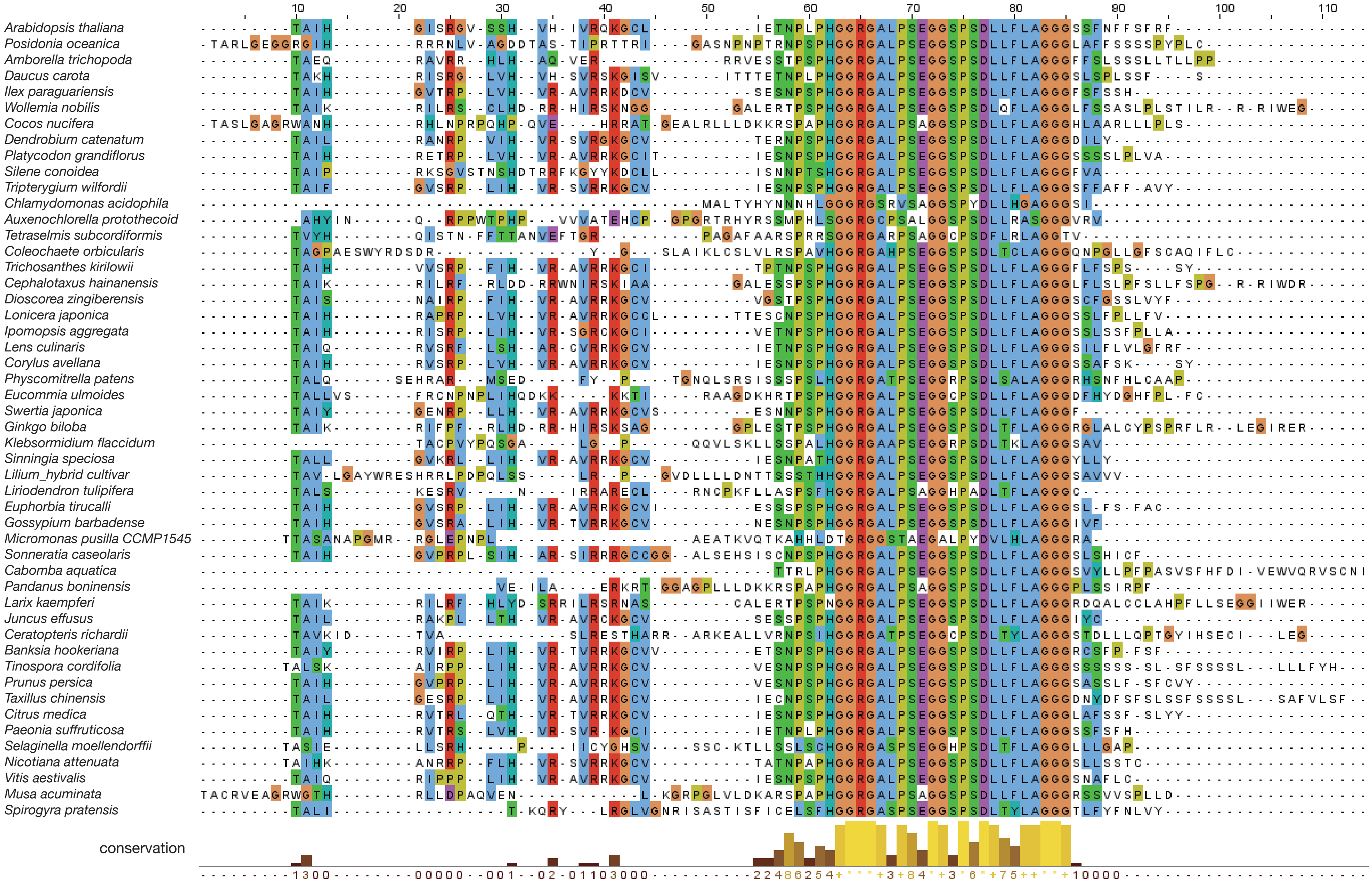

#### AT4G34710 *SPE2*

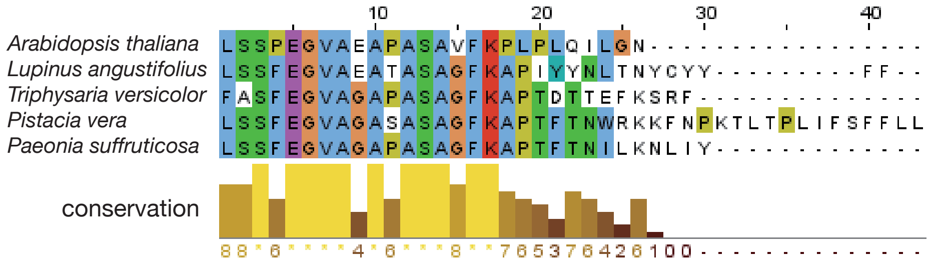


#### AT4G38520 *APD6*

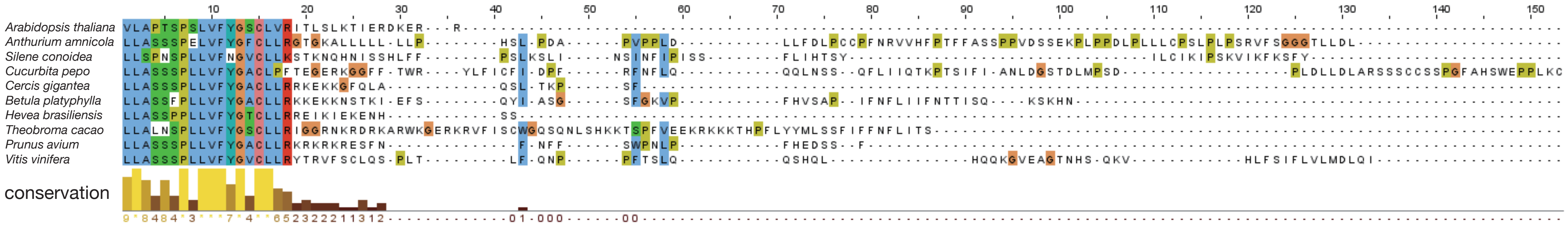


#### AT5G35750 *AHK2*

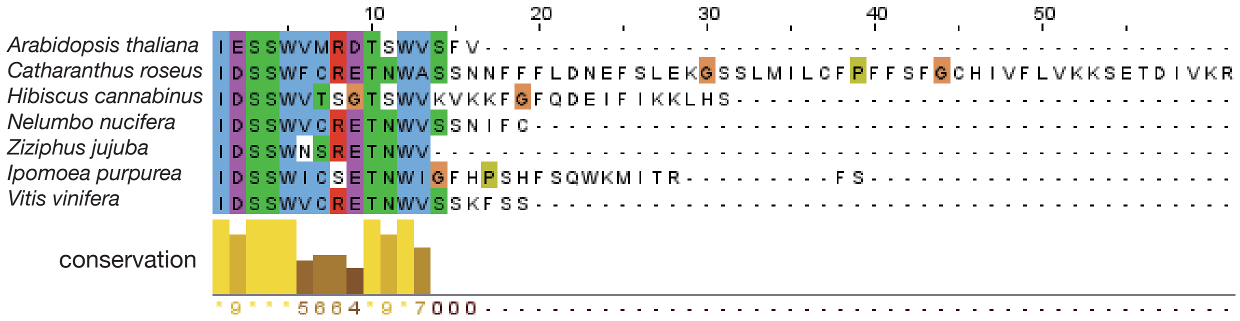

### AT5G55120 *VTC5*

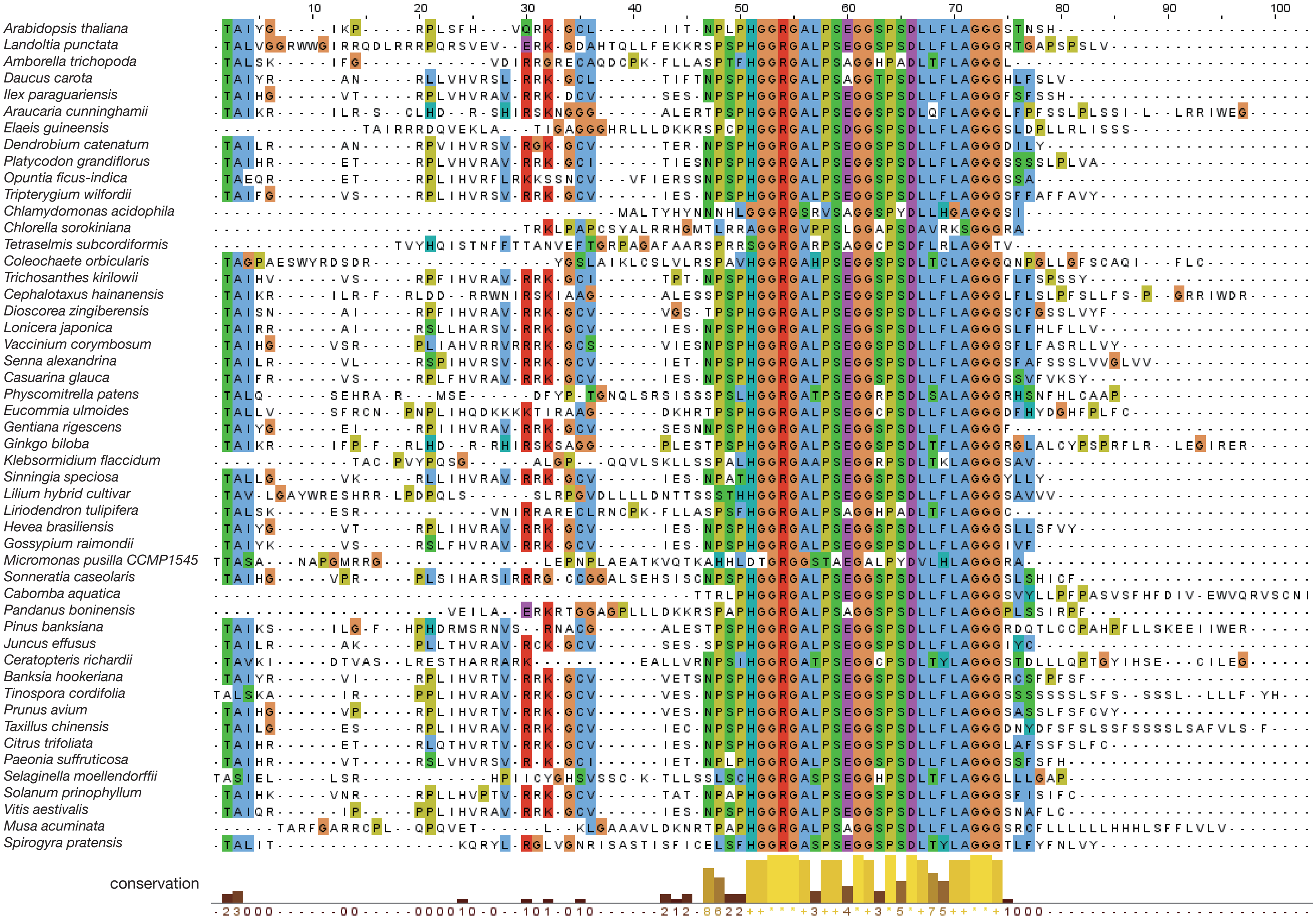

**Supplementary Figure S2.** Alignments of the novel and known conserved non-AUG uORFs identified in this study. The amino acid sequences of the Arabidopsis conserved non-AUG uORFs and their homologous uORFs were aligned using Clustal Omega version 1.2.2 and displayed using Jalview version 2.10.2. For each alignment, representative uORF sequences were selected from orders in which uORF-mORF homologs were identified.

### Supplementary Figure S3

**A** AT1G32240 (*KAN2*)

```
                                                                    A
  1 AGCCCCCCATGTCCACACAAACACCTTCTCTCTCCTCTCTCTCACACACACTCAAGTGTC
                                                         V  S

 61 ATTTTCTCACATATACTTCACACAAAAGAAACAACCATTCACTACATACTACCTTGTCAT
     F  S  H  I  Y  F  T  Q  K  K  Q  P  F  T  T  Y  Y  L  V  I

121 TGATTTTCTCTCTCCCTCTATATCTCTCAAAAATCTTTGCAAAATCTTCTTAACCTCAAA
     D  F  L  S  P  S  I  S  L  K  N  L  C  K  I  F  L  T  S  N

181 CAATATAAACAACATTCTCAACAAAACCCAGAAATCTCTTCCTCAAATCCTCAATACCCT
     N  I  N  N  I  L  N  K  T  Q  K  S  L  P  Q  I  L  N  T  L

241 TTTCTATTTTCCTCCCTATACCCTTAATAAAAAAGGATATCTTTCTCTTTTTATCTTCTT
     F  Y  F  P  P  Y  T  L  N  K  K  G  Y  L  S  L  F  I  F  F

301 TGTCCTAATTCAAGAAAACAAGAACTAGGGTTCATCATCTGTGGAAACCGACTGAAGAAT
     V  L  I  Q  E  N  K  N  *

361 CTCTTCAAGAAGAAGTTAAAGGTCTCTTTCATGGA
```

**B** AT1G53300 (*TTL1*)

```
  1 GGTTCTCCATCTTCACTCCTCTTCTTCTGTGAACCTTCTCTCCTCTCTCTCTTCTCTTCA
                                                  G
 61 CTGAGAAAAAAACAAGTTTTCTCTCTCTAGGCACTCACATTCATTCTTTGCTTCAACCTTT
                                               I  L  C  F  N  L  L

121 TAATTCACGAGATCCCCAATCTCTCTTTCGATGCCTTTTCACAGAGTATCTAATCCTCCT
      I  H  E  I  P  N  L  S  F  D  A  F  S  Q  S  I  *

181 CCTCCTCCATTCATCTTCTCCTTCTCCATTCTTCTCCGTCGTCTCCATTAACAAACAACC

241 ATTTCTCTCACCGGAACTCCATTTTAGGGTTTTCTTCGATTCTCTCAAATTTCATTCCTT

301 TCACTCACCACAACACTCAAAATGCC
```

**C** AT1G58290 (*HEMA1*)

```
                                                 AA
  1 ATAAATACTTATATGTTGCCGTGTAAGAACAAATGCCACCAAATACGCAAAAACGAGTGT
                                                               V
                                                               V
    C    A
 61 GGATAAAACGTGTTCTCCACCCGCCAATCAAATTCCATCAAAATCTCAATGCATTCCTCA
     D  K  T  C  S  P  P  A  N  Q  I  P  S  K  S  Q  C  I  P  H
     D  KN  V  F  S  T  R  Q  S  N  S  I  K   S  Q  C  I  P  H

121 TTACATATCTTTAGATTACCATCCAATCATAATCCTCAATTTCCAGTTCTACTAAACCGG
     Y  I  S  L  D  Y  H  P  I  I  I  L  N  F  Q  F  Y  *
     Y  I  S  L  D  Y  H  P  I  I  I  L  N  F  Q  F  Y  *

181 TTTTGACCCGATTTGATTTTTCAGCCTCAAATCTGACAAATCATATTCACAGCGATCTCC

241 TTCTTCCACATCGCAACCCAATCCCTCTCTGTCTCTTCCTCAATTCTTATTCTCTCTCGC

301 AAAGGTTTGGTTTTTTGAAGACAGAGAGGAGATTTGAGATTTTGGGTTTCAATGGCGGT
```

**D** AT2G16720 (*MYB7*)

```
  1 ATAATGGTTATGAGATCTAAAACATCAAAGGAAAATCAAGAAGAGAAAGGTATAAAATAT

 61 TGAATAAATAATATGAATGTGGCAGTTATAGTTTGATGTAGAATAAAAGATTAAACAAAA
                                      C
121 ATCAAGAGCCCCACCAGCCACTCCTTCAAACGTGTCCTACCAACTCTCTTCTAGAAGCAA
                                   V  S  Y  Q  L  S  S  R  S  N

181 TTAGGCTCACCTTCCACCTCCAACACCTCTCTCCCCTTCTCTCTCCATAAAACCCTCAAC
     *

241 AACAGAATCAAAAGCATCATCATCATCAGAAGAAGAGAGTCATGGG
```

**E** AT2G20080 (*SPEAR1*)

**C**

1 GGCGTATCTATTGAA**ATCTGGAAACAATCTGATCCAGCGATGCAGTAG**TTATACACTAAG

**I W K Q S D P A M Q ***

61 GTTTTCTATAAAAACTCAAAATGAATTGTAAAAAAGATTATAAAAAAGAGATCACCAATT

121 CACCATGAAAGAGATCTCTCCAATTTCTTAGCTCAAAGCAAAATAAGAAAATTCATTTCT

181 TCTCTCTCTCTCCCTCTCTTTCTCTCTAGAGAAGACTTTCTCTCTCTAGTTTCTTGAATT

241 TTCCGGTATACCTTACGATCGCAAGTAAAATTTAAGGCCGATCAAAGAAACATGGG

**F** AT3G05470 (*FH11*)

1 GCCAACAATAAGAGGACCCACTTCACACCAACACCTTCACGCTTCTTTCTATCTTCTCTT

61 TGCTTACTAGATTTTCTCCTTTTTAATCTCTTCCTTAAACAAAAAAGCTGTTTTCAAACT

121 TAAACTAAAAAGCTTCATATTGTCACCACATTAATTCCAAGAAAATTCGCATTAATGCTC

181 TGCAAAATTTAATGATTTTAAAATTAACATTTGTTCCTTTATTCAAGAAAAATTCGCATT

241 AATTTATTTATTTAAGAGATGATTACCTAAAATGGTGTGTATAAATTGCAAAATTAGCTC

301 TGTTGTTCTGCTAGGAGAGTTAATGGAGAAAGAGGGTGTTACCATTACCAGTACAACTAT

**A**

361 AG**TTGCAGAGTTCAGGGGTTCCTTGGAGTAGCATTAAAGTCTCAAAGGAGAGAAAAATAA**

**L Q S S G V P W S S I K V S K E R K I K**

421 **AAAAAAAATACATTAAAGACAAATTGAGTCTTTTCTCATAA**ATGGTTTCTTTGTAGTGTT

**K K Y I K D K L S L F S ***

481 TATGATTTGTTTTTACAGAGAAAAGATCTAAAAAGAGAACAAAGATATGGT

**G** AT33G08850 (*RAPTOR1B*)

```
  1 GAAGTGGAAAGAAAACGCAAATGCATTGAAGACTTTCCCTTTGTCTCTCTCTTTGACAAA

 61 ATCCCAATTTTCGCATCCTCACTTTCTCCTCTGCCTTTTTTTTTTCTCTTCTGCCGCCAC

121 CACCACCACCACTGTTAATTCGCCGCCGCTTCCACATCAATCGCCGTCGTAGCCAATCAT

181 CATCGTTCTTCATCCTCCGCTTCTCCTGATTCTTAGGGTTTCCGCTTCCGCTTCTCTGTC
                     CC                    G
241 GAGGGTTTTTCGAGCTGGATTCTCTGCTCCTGCGCCGCTGGTGCAGCGGCTTTGTTGATT
                  L  D  S  L  L  L  R  R  W  C  S  G  F  V  D  *
                  L  D  S  L  L  L  R A  L  V  Q  R  L  C   D  *

301 GATCTGGATTTCTCCGTCTTCCTTTCTATTTTATTTTCTGTATCTCGCACAATCCAAACA

361 AAAAAGCTAGGGCTGAGGATTTGATTTGCGTGATTTCTGGGGGTGTGTTAATTCGGGGGT

421 GTGTTCTCCGATTTGGATGGC
```

**H** AT4G03260 (*MASP1*)

```
  1 GCGTTGCTTTGCCTCAAACTCAAATTTTGAAGATAATTTTTTTGGTTTCTCTGCTACTGA

 61 TTTCGAATCTTTACGCTTAATCAATTATTCATCTGGATTTTCAAATCCAATCCTTTTCCT

121 CCCTGGTGACCAAGTAATTGGGGATTAACAAAGGGACTGGGGCTTTATTAAGCTCGAAAC
                      C
181 AATTGTTCTTCACTCTGGCCGCTTCTGCAAAGTCGTGTGCTTCTCGCACCTCTGCGCAGG
                  L  A  A  S  A  K  S  C  A  S  R  T  S  A  Q  V
                  L  A  A  S  A  K  S  C  A  S  R  T  S  A  Q  V

241 TGTGGGGTGGCCTTGAAGTGGTGACTTCACTCTGTCTTCCAAGGATTTCTCTAGCTTCTA
      W  G  G  L  E  V  V  T  S  L  C  L  P  R  I  S  L  A  S  I
      W  G  G  L  E  V  V  T  S  L  C  L  P  R  I  S  L  A  S  I
                                                        T
301 TCATTTGGAGGCTCCGTCGTGTTTCTGAGGGTCTTACCCTTGATTACGGCTAACATTTAC
      I  W  R  L  R  R  V  S  E  G  L  T  L  D  Y  G  *
      I   G  G  S  V  V  F  L  R  V  L  P  L  I  T  A *

361 TACGCCCACTATACCACCTCTTAAGCTTTTATAAGAACTTTTTATTCGTGCAAAAACTTA

421 ACTTGGATCTTAATTGTTTCTAGTTTAGTTGTGGTTGTAAAGTAAAGAACCGTTTTTTTG

481 TGGTTTGGTTTTGCTTTTTTAGTTTCCGTTGGTGATGTAGTTCGATGTTATTAAGTTTGA

541 CACAATGGTCAGGTT
```

AT4G19110 (*MPK22*)

1 ACGAGAGAACCGCACATTGAAAAGGGAAGGAAGAGAGATATTTGGCAGAGAGAGAAAACG

61 TTGGAGGAAGGATTGATATTTGTTGGACTTTGGATCGAAAAATTGAAAGGTAAAAGAGGT

**C**

121 AGAGAAGCT**CTGGATTCCGTCTCTTCCCGTCGTCGGCGACGGTGGTTTTGCCCGTTGGCT**

**L D S V S S R R R R R W F C P L A**

181 **CTGAGGTTATATCAGGCTTTAATTTAA**GTGGATTGTCATCTTCTGAGAT**ATGGAACAAGT**

**L R L Y Q A L I * M E Q V**

241 **TTTTGTCTGGCCAAGCTGCTACCACTATCGGCTTTTCTCATTCCAAGAAGCGCTCGATTG**

**F V W P S C Y H Y R L F S F Q E A L D W**

301 **GCGGTTTCTTGTACGTTCTGATTTCCTTGTGGGCTCTTTCGTTAACTGCACTTAG**TGGTC

**R F L V R S D F L V G S F V N C T ***

361 AAACCCTCTTCTATGATCTTTAGGGGAAGGTGTGAGGCAAATTTCTGAGACAGGGCTGAA

421 GTCTGCAAATTCTTCCTTCGCATAGTCCTAGAGAAACCTCTTTGTTCCTCTCTTTCATAT

481 CTTGTTCTTATGGGTGCTGGTTGAAGAAAGATACATCAATTTCTTTTCTTTTTCAGTTTA

541 CAAATGTCGTTTCCTTGAGTCTGAACCCCTGAGTGAGATGTTGTGAGCATAGGGTCCTTC

601 CTATTAGCCAAAACAGTCTTTCTTGTTGGTTCCTTCGCGTCTTTGAGTTCTCAGTATGTG

661 TTTTCCTCTTCTCTTGCTCTCTTTCTTTTGCCTAAATGGTCAACCCGTTTATGTTCGAAG

721 TAAATAACGGTCAGTCTACCCAGCTGGAGAATCATTGATTTTACTGCCTTGGTGTCAAGT

781 ACTTCCTACGAACGACTGGAAGATTAATGCTATATAAATCTTTCAGAAGTTCTTTACTTA

841 CATAATATAAAGAGGCTACTTTGCTTAACCAGAAAAAAAAAGATGGA

**J** AT4G34710 (*SPE2*)

1 ACGCCTTCTCTTCTTCTTCTTCTTCATTTCCGTCTTTTCCGCTAGGTTTCCGTCTCTGTT

61 TTTTCTTGCATTTTCCCCAGAATTCCTTCACATTCTCTCCTTTTCCTCTTTTTAATTCCT

121 CACCACCTCAAATTCTTCATAAACAAATCCACTTCTGCATGGTTGGGAGAATCTGTTAGA

181 GAATTTTTGTTGTTGTTGGGTTCAGATTTGGAAAATTTCCTTTTTTTCGAGTTTCTTCTG

241 AATTATTCATCGTGGAGATTCTCGGATCGGGAAGGGCTGCGAGAGGTGGTTTTCACGGCG

G C T C

301 GGGATAGATTCATCTT**CTGTCTTCTCCTGAGGGGGTAGCCGAGGCTCCGGCCTCGGCGGT**

**L S S P E G V A E A P A S A V**

**L S S P S G G S R G S G L G G**

361 **TTTTAAACCCCTACCTTTACAAATTCTGGGAAATTAG**AAAAAAAAAGGTTTGATTTTTCA

**F K P L P L Q I L G N ***

**F K P L P L Q I L G N ***

421 GTTTTTGCTCTGTTCTTCTCTCGAAAGTTTTTGTTTTGTTCGAGAATTTCTGTAAAGTGA

481 GATTGTTGATCGTTTGCTGATAAGAGGGTGAAGATAAAGATGCC

**K** AT4G38520 (*APD6*)

```
                                               C
  1 TGGAGCCCGAAAGCTGGGACCTTTGATGTGCTGGCTCCGACCTCGCCATCGTTGGTCTTT
                               V  L  A  P  T  S  P  S  L  V  F

 61 TATGGGTCTTGTCTGGTAAGAATCACTCTTAGTCTTAAGACTATAGAGAGAGATAAAGAG
    Y  G  S  C  L  V  R  I  T  L  S  L  K  T  I  E  R  D  K  E

121 AGAAGATAAAAAAAACAACTCCAACTTTCTTTCTTTTTTTTACATAAAAACGTCGAAGGA
    R  R  *

181 GGATGTCTCGTTCGGCGACTTGACCTCTCTCTCTCTCTCTCTTTTACCTTCATTGTTCAC

241 TGTTTCCAGACATTGAAAGAAAGATCATTAGTTTCTGAGGGGATAACCATTTTTTTTTCT

301 TCTTAAAAAGTTTTTTTTTTTGTCAATTGGTCTTCTTCTTCCTCATCCGCTCGCTTATAT

361 TTAATTTAATAATAATAAAAATCGGTGTTCATTTTTATATAAAAGTATAAATTTTGTGTT

421 TGGTTGATCCATCCGTAGTCGTAGTAGATCTACAAGCTCTGAAATTCCACGCCCACATCT

481 CCTCCGTCCTCTTGAGATCCTTCTCTTCATTCATTTTCTTGCTCAACTTGAGTTGCTTCC

541 AACATCTTGCAAGATTGTCTTTGAGTTCATTCAGGTCTTGAAGAAGAGCACTCTTTTAAG

601 CTGTGTGAAACTTGCGCCGTTCCTGTGAAGAACTACTTTTGGCATTGCAGATTTCAGAGA

661 GCTGTGATTTGTGCTATCTTAAAAACGGACAAGTTCTATGTTGTGGCGGATGAGATGAGG

721 GATGCT
```

**L** AT5G35750 (*AHK2*)

G

1 ATAATTTCTCTAACAAATGCTTCTTTTATCCAAACTCTGATC**ATTGAGTCCAGTTGGGTA**
I I S L T N A S F I Q T L I **I E S S W V**

61 **ATGAGAGACACCAGCTGGGTTTCTTTTGTTTAA**GCTCACAAACATCTAAAACAGAGAAAG
**M R D T S W V S F V ***

121 CTTTGTCATTTTTCTTTACCCTGAAACCATGAAAGAAGATAAAGCTTTGTCCTTTTTCTC

181 TTTTCCGTTTCTTTACTCTCAACAATGTCTTCTTTCTCCCATTGCAACAACCACTGGGTT

241 TGGTTCATCTTGTCCTCTTGCGTCTTAAGTAAGAAAAAAGAAAAAAGCTTTGCGTTTGGG

301 ACTATGTCACAGCTTCTTGTTCTTCTTGTGGATGGTTTCAAGAACTCAACTATGAGATTG

361 TTGTTGATAGAGTTTTAGTTTTGGGTTTGAAGAGATCTGAATTTCTTCAGCTGAATCTGA

421 GATTAGGAGTCGAAATGTC

**M** *Theobroma cacao MYB7*

C

1 TATCATCAAGACACAGGCCACCAGCCACTCCCTCAAAC**CTGTCTCACCAACTCACATCTA**
**L S H Q L T S R**

61 **GAAGAAAATTAGGGTCACCACCCCTTTTCTCTCTCTTTCCCTCCCTGTGTGTGAGAGTAA**
**R K L G S P P L F S L F P S L C V R V I**

121 **TAGTCAAAAGGCTATATAACCGCATCTAA**AGTTCTCTCAAAGGGCCGGCCTCAATGTTCC
**V K R L Y N R I ***

181 AGCAACGCAAAAAACAAGGCCTCCATTCCATTTTCCACCACCACTATTCTACTTCTTCTA

241 AATTCTAAACATATAAATTTATACTACTTACTGCTACTACTATTAAAAAAAATAACACAG

301 CAACGTAGCTACTACTGCTAGTTTCCACAACTTGGAAAAGTTTCTAGTACTCAAATGGG

**N** *Arachis ipaensis FH11*

1 ACAGTGTTATGTCCATGCATTAAGTTTGTCTTCAAAATTATTAAATGTGAGAAGAGATAA

**C**

61 GAGTGTATGAACAGTAAAGAAAAGAAAGTCCCAATTTACAACCACAA**CTGCCAGGTTCAG**

**L P G S G**

121 **GGGTTCCTTGGAGTAGCATTAAAGACACTGTTTTCAGTTTCAGATCTTCTATTGCACACA**

**V P W S S I K D T V F S F R S S I A H I**

181 **TTACATACATTACCAGCACCAAAAGAAAAAACCCCAAAAAAGAAATGGAACAACAAAAGG**

**T Y I T S T K R K N P K K E M E Q Q K G**

241 **GTAAAAGACAAGAAGCTTTTATAAGGGGAAAGCTGCATTGA**TTTCTAGATGACATTTTAG

**K R Q E A F I R G K L H ***

301 TTGGAATCAGAAGAAGTAAGAAGAAAGAAGAAATAATGGG

**O** *Theobroma cacao APD6*

**C**

1 CCATAGTGGTGTGGAGCCCAAAACTGGACCTTTGCT**TTGCTGGCTTTAAACTCTCCATTG**

**L L A L N S P L**

61 **TTGGTCTTTTATGGGTCTTGTCTGCTGAGAATTGGAGGGAGAAACAAGAGAGATAGAAAA**

**L V F Y G S C L L R I G G R N K R D R K**

121 **GCAAGATGGAAAGGAGAGAGAAAGAGAGTTTTTATAAGTTGTTGGGGCCAAAGCCAAAAT**

**A R W K G E R K R V F I S C W G Q S Q N**

181 **CTTTCACATAAAAAGACTTCTCCTTTTGTTGAAGAAAAAAGAAAAAAAAAAACTCATCCA**

**L S H K K T S P F V E E K R K K K T H P**

241 **TTTCTCTATTATATGCTTTCCTCCTTCATCTTCTTCAATTTTCTCATCACTAGTTGA**AGC

**F L Y Y M L S S F I F F N F L I T S ***

301 CAACCAAATTCCAAACTTTGTTTTTTCTCTTGCTTTGATTTGATTGATCCATAGATCTGC

361 TAATTCCAATTTCATGCCCAGATGAAGTTTTTTCCTTTCCTATTTCCAGATCCTCCTTTT

421 CAATTATAAACCAAAGGGGTCAAATAGAAAGTGTACAATTATTGTGTTGAGAAGCAGAGG

481 ATACGGTTCACTGTTCAGACTTCTAGGGTATAGTTTATATTCAGTGGTTATTTAGATGTG

541 GAGAATTAGGAGGGGTTTTTGGATGAGATGAG

**Supplementary Figure S3.** Nucleotide sequences of the 5′ -UTRs used for the transient expression study and the deduced amino acid sequences of the conserved non-AUG uORFs. The 5′ -UTR nucleotide sequences are based on NCBI RefSeq transcripts [GenBank accession numbers: NM_102957.4 (A), NM_104208.3 (B), NM_104609.4 (C), NM_127224 (D), NM_201757.2 (E), NM_111420.2 (F), NM_111719.3 (G), NM_116564.4 (H), NM_179076.1 (I), NM_119637.3 (J), NM_120013.6 (K), NM_122966.3 (L), XM_007041801.2 (M), XM_016321042.2 (N), and XM_007041714.2 (O)] and Plant Promoter Database version 3.0 (http://ppdb.agr.gifu-u.ac.jp/ppdb/cgi-bin/index.cgi). The nucleotide sequences and the deduced amino sequences of the conserved non-AUG uORFs are shown in bold. The replaced nucleotides in the NCC and AUG mutants were shown in black background. The AUG initiation codon of the main ORF in each sequence is boxed. Arrows indicate the positions of the primers used to clone the 5′-UTRs. The nucleotide sequences of AUG uORFs are underlined. In (C), (G), (H) and (J), the nucleotides that were deleted or inserted in the frameshift mutants are shaded, and the deduced amino sequences of the frameshift mutant versions of non-AUG uORFs are indicated. In (J), the nucleotide ‘G’ replaced by ‘C’ in the frameshift mutant is shaded and underlined.

### Supplementary Figure S4

#### AT4G34710 (*SPE2*)

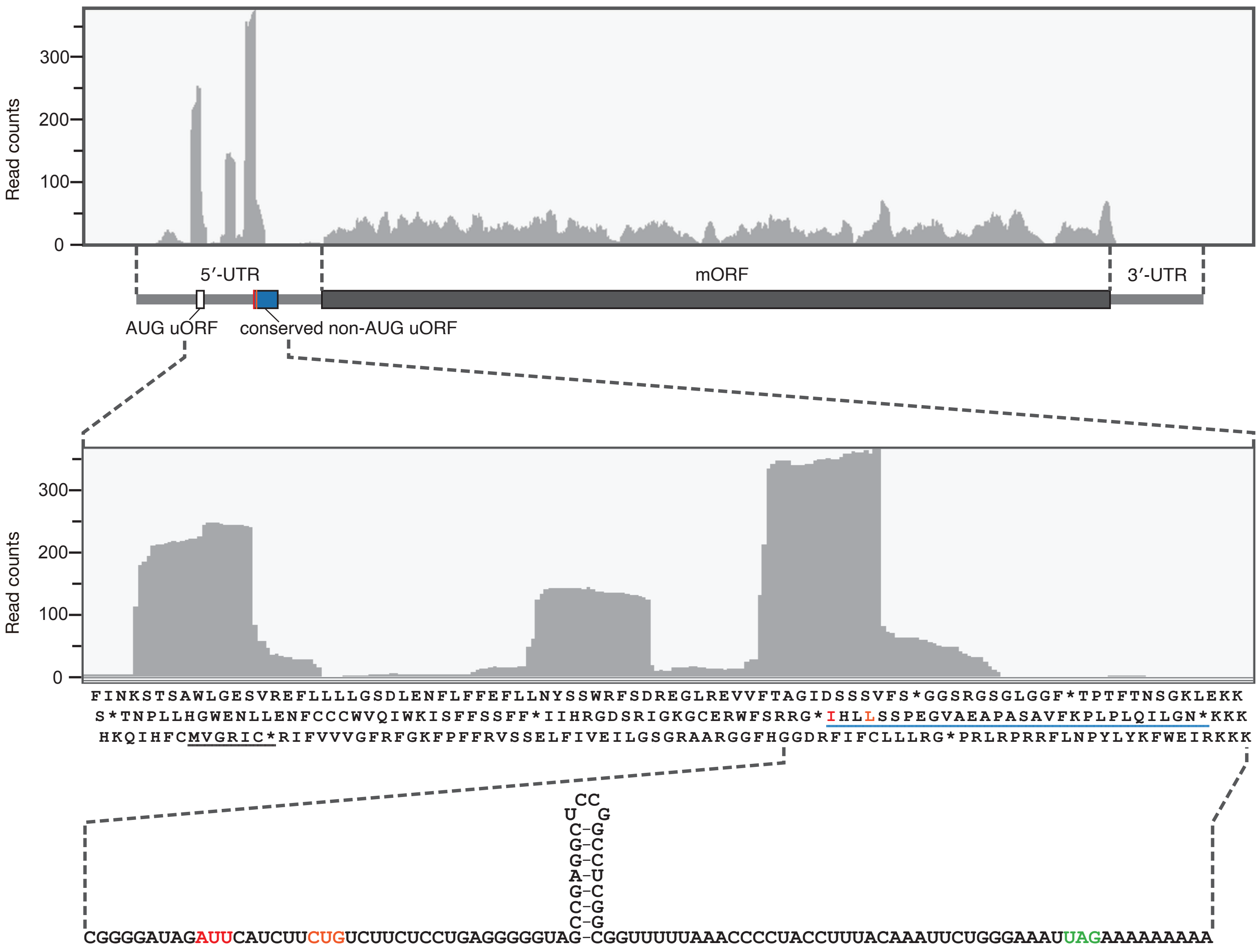


**Supplementary Figure S4.** Ribosome footprint distribution along the *SPE2* mRNA. The *A. thaliana* ribosome profiling data from Kurihara et al. (2018) (SRA accession SRR6466829) was analyzed. After removal of the adapter sequences, low-quality reads and reads derived from rRNAs and other non-coding RNAs were discarded. The 27-29 nt reads from the resulting data were mapped to *A. thaliana* reference transcript sequences (TAIR10). Ribosome footprint (read) distributions along the entire *SPE2* mRNA and a part of the 5′-UTR are shown in the upper and middle panels, respectively. The red and orange lines in the blue box indicate the positions of the AUU and CUG start codons of the conserved non-AUG uORF. In the middle panel, the deduced amino acid sequences of the three reading frames of the corresponding region are indicated. The amino acid sequences of the AUG uORF and the conserved non-AUG uORF are underlined in black and blue, respectively. In the bottom panel, the hairpin structure in the conserved non-AUG uORF is shown, which was predicted by CentroidFold (http://rtools.cbrc.jp/centroidfold/). The AUU and CUG start codons and the UAG stop codon of the conserved non-AUG uORF are highlighted in red, orange, and green, respectively.
